# The SARS-CoV-2 nucleocapsid protein associates with the replication organelles before viral assembly at the Golgi/ERGIC and lysosome-mediated egress

**DOI:** 10.1101/2021.06.15.448497

**Authors:** Katharina M. Scherer, Luca Mascheroni, George W. Carnell, Lucia C. S. Wunderlich, Stanislaw Makarchuk, Marius Brockhoff, Ioanna Mela, Ana Fernandez-Villegas, Max Barysevich, Hazel Stewart, Maria Suau Sans, Charlotte L. George, Jacob R. Lamb, Gabriele S. Kaminski-Schierle, Jonathan L. Heeney, Clemens F. Kaminski

## Abstract

Despite being the target of extensive research efforts due to the COVID-19 pandemic, relatively little is known about the dynamics of SARS-CoV-2 replication within cells. We investigate and characterise the tightly orchestrated sequence of events during different stages of the infection cycle by visualising the spatiotemporal dynamics of the four structural proteins of SARS-CoV-2 at high resolution. The nucleoprotein is expressed first and accumulates around folded ER membranes in convoluted layers that connect to viral RNA replication foci. We find that of the three transmembrane proteins, the membrane protein appears at the Golgi apparatus/ERGIC before the spike and envelope proteins. Relocation of the lysosome marker LAMP1 towards the assembly compartment and its detection in transport vesicles of viral proteins confirm an important role of lysosomes in SARS-CoV-2 egress. These data provide new insights into the spatiotemporal regulation of SARS-CoV-2 assembly, and refine current understanding of SARS-CoV-2 replication.

## 1 Introduction

SARS-CoV-2 is an RNA virus and the causative agent of COVID-19 (*1*). To date more than 176 million cases of this disease have been diagnosed, resulting in more than 3.8 million deaths (*2*). Great efforts have been made in the development of measures for containing the spread of SARS-CoV-2, including the repurposing of previously produced drugs (*3*), therapies (*4*) and the development of vaccines (*5*).

While new diagnosis, prevention and treatment options for COVID-19 continue to emerge at a rapid pace, the understanding of the biology of SARS-CoV-2 advances more slowly. Unravelling the mechanisms of transmission and replication of this virus is crucial for the development of rationally designed drugs and vaccines, and to understand the long-term effects of the disease, allowing researchers to develop countermeasures against evolving SARS-CoV-2 variants of concern.

SARS-CoV-2 spreads among humans primarily via respiratory droplets when two individuals are in close proximity (*6*). It is an enveloped virus that enters the cells of the respiratory tract through the interaction of the receptor-binding domain on the spike protein and the angiotensin-converting enzyme 2 (ACE-2) receptor on the cell surface (*7*). The positive sense, single-stranded RNA genome of SARS-CoV-2 is then released into the host cell cytosol and is directly translated. Two large open reading frames (ORF1a, ORF1ab) are translated into large polyprotein complexes (pp1a, pp1ab), which are co-translationally and post-translationally cleaved to generate 16 nonstructural proteins (nsp), for which characterisation is ongoing (*8*). The remaining ORFs encode the four structural proteins of SARS-CoV-2 (*9*). In coronaviruses in general, the nucleocapsid protein encapsulates the viral RNA (*9, 10*), the spike protein mediates cell entry (*7*), the membrane protein is embedded in the envelope and thought to provide a scaffold for viral assembly (*11*), and the envelope protein forms ion-conductive channels in the lipid viral envelope (*12*). Upon infection by SARS-CoV-2, the virus initiates the biogenesis of replication organelles (ROs) containing interconnected perinuclear double-membrane structures such as double-membrane vesicles (DMVs), which are derived from, and tethered to, the endoplasmic reticulum (*13*). It is assumed that these structures protect the viral RNA from degradation by cellular RNAses during genome replication (*14*). This hypothesis has been corroborated by the recent finding of Klein et al., who showed the presence of viral RNA in the DMVs (*15, 16*). The DMVs possess a pore in their double membrane lining, by which the RNA is thought to access the cytosol (*17*). The assembly of mature SARS-CoV-2 virions occurs within the endoplasmic reticulum to Golgi intermediate compartment (ERGIC) (*8*),(*13*). The egress of coronaviruses is assumed to occur via exocytosis (*15*). Recent evidence suggests that newly formed SARS-CoV-2 virions reach the cell periphery using lysosome trafficking (*18*).

The interactions of each SARS-CoV-2 protein with a series of host cell proteins have been partially studied by combining light microscopy and proteomics (*19, 20*). Gordon et al. (*19*) used confocal microscopy to study the distribution of two of the structural proteins of SARS-CoV-2 in infected Caco-2 cells at one time point post infection. The imaging revealed a cytosolic signal for the nucleocapsid protein and strong interaction of the membrane protein with the Golgi apparatus. In the current work, we provide more detail on the interplay between all four structural proteins of SARS-CoV-2 and the host cell. We furthermore image and analyse multiple time points over the time course of the infection cycle.

The replication of SARS-CoV-2 is known to extensively change the localization and reshape the morphology of cell organelles and the cytoskeleton within the host cell. Such morphological alterations have recently been studied in Calu-3 cells using both optical and electron microscopy (*13*). The study by Cortese et al. analysed infected cells at a series of time points post-infection to detail the progression of the viral cycle, focussing on the host cell structures in detail. They demonstrated the progressive fragmentation of the Golgi apparatus, the recruitment of peroxisomes to the sites of viral replication, and the reshaping of the vimentin network to accommodate the DMVs. While electron microscopy highlighted cellular structures with high definition, the viral proteins were visualised with lower resolution using confocal microscopy. The power of super resolution optical microscopy has been demonstrated by application of 3D-STED (stimulated depleted emission microscopy) to reveal the interaction between double-stranded viral RNA and the vimentin network. However, the latter technique cannot be performed in high-throughput fashion and is not easily adapted for multiplexed imaging of several proteins simultaneously.

Here, we employ a range of light microscopy techniques to overcome some of these limitations. We present a detailed investigation of the spatiotemporal organisation within the host cell of the structural proteins of SARS-CoV-2 during an infection cycle. Specifically, we focus on the assembly of these proteins to form mature virions and their interaction with host cell organelles that are affected by SARS-CoV-2 infection. In order to obtain high quality imaging data, we used Vero cells for infection since their morphology is well suited for fluorescence imaging. In addition, numerous SARS-CoV-2 studies based on Vero cells exist allowing us to put our results into context. We present a fixation protocol that permits transport of infected Vero cells from class 3 containment laboratories to high resolution imaging facilities. Establishing immunostaining protocols for the imaging of multiple SARS-CoV-2 proteins simultaneously provided well defined and controlled snapshots at different infection stages. In order to achieve this, we employed a combination of widefield, confocal, light-sheet, and expansion microscopy. Imaging was possible in up to four colours at subwavelength resolution, providing details on the intracellular trafficking of all SARS-CoV-2 structural proteins in space and time during the course of the infection. By combining expansion microscopy and light-sheet microscopy, we have produced volumetric maps of protein distributions in whole infected cells. We furthermore visualise SARS-CoV-2 induced morphological changes of host cell structures that are involved in assembly and egress. We find that reshaping of microtubules, relocation of lysosomes and fragmentation of the Golgi apparatus largely correlate with the local accumulation of the three viral transmembrane proteins spike, envelope and membrane protein.

## 2 Results

### 2.1 The cellular distribution of SARS-CoV-2 structural proteins is tightly regulated in space and in time

We first optimised cellular fixation and staining protocols, using transfection to express the four structural proteins of SARS-CoV-2 individually in Vero cells. We also optimised fixation (formaldehyde and glyoxal) and permeabilization (Triton X-100 and saponin) reagents. Each cellular structure has its own ideal immunostaining conditions (Supporting Figure 1); the endoplasmic reticulum was best fixed in a glyoxal buffer, preserving the fine structure of the tubular regions. In contrast, the Golgi apparatus was only stained when fixed with formaldehyde independently of the detergent. As a final example, lysosomal staining was only achieved after permeabilization with saponin. The optimal staining conditions for the cellular structures being investigated determined the choice of experimental conditions for each sample. A summary of the optimised fixation and permeabilization conditions for each of the structures investigated in this work is presented in Supporting Table 1.

The immunostaining of transfected cells with the selected antibodies was successful in all fixation and permeabilization conditions tested (Supporting Figure 2). Interestingly, the pattern of the spike (S) protein staining was different in the two fixation conditions (formaldehyde and glyoxal) tested in transfected cells. We did not note any differences when fixing and staining infected cells in these two conditions. It has been previously observed that the intracellular localization of viral proteins can significantly differ when comparing an individually expressed viral protein and the same protein within an infected cell (*19*). These observations confirmed that investigations should be carried out in virus infected cells.

In this study, we fixed infected cells at multiple time points post-infection (5, 7.5, 10, 12 and 24 hours). Infection stage varied between individual cells in the population. The spatial distribution of the viral proteins changed over time and varied between individual cells at the same time post-infection, particularly at later time points. This is expected since experimental conditions such as the multiplicity of infection (MOI) influence replication kinetics. However, we identified similarities in the expression and distribution of the viral proteins in individual cells within the heterogeneous population and across time points. These patterns correspond to distinct events in the replication cycle. We used these patterns to classify the cells into different categories. This analysis on single cells rather than the population average proves useful as it allows us to gain a clearer picture of how the virus cycle is staged in time as well as to connect certain steps in the viral replication cycle with morphological changes in the host cell.

We found that three different categories or stages were sufficient to classify the status of any cell in the population (Figure 1A). At an early infection stage (5 hpi), cells were seen to express the nucleocapsid (N) protein only. At this stage (referred to as stage 1), the N protein is not homogeneously distributed inside the host cell cytosol, but forms small puncta. From 7.5 hpi onwards, the other structural proteins of SARS-CoV-2, S, membrane (M) and envelope (E) protein, are expressed and localise in a compact juxtanuclear membrane compartment (stage 2). Cells which were characterised by fragmentation and spreading of the compartments containing the S, M and E proteins were classified as stage 3; in these cells, the N protein is homogeneously distributed in the cytoplasm. At stage 3, we observed small dots of all four structural viral proteins at the plasma membrane, indicating the formation and trafficking of mature SARS-CoV-2 virions. These relative timings within the replication cycle were not previously known in this detail. The findings were enabled by the classification strategy described here, which considers the staging of cells individually rather than population averages at different times post-infection.

**Figure 1:**
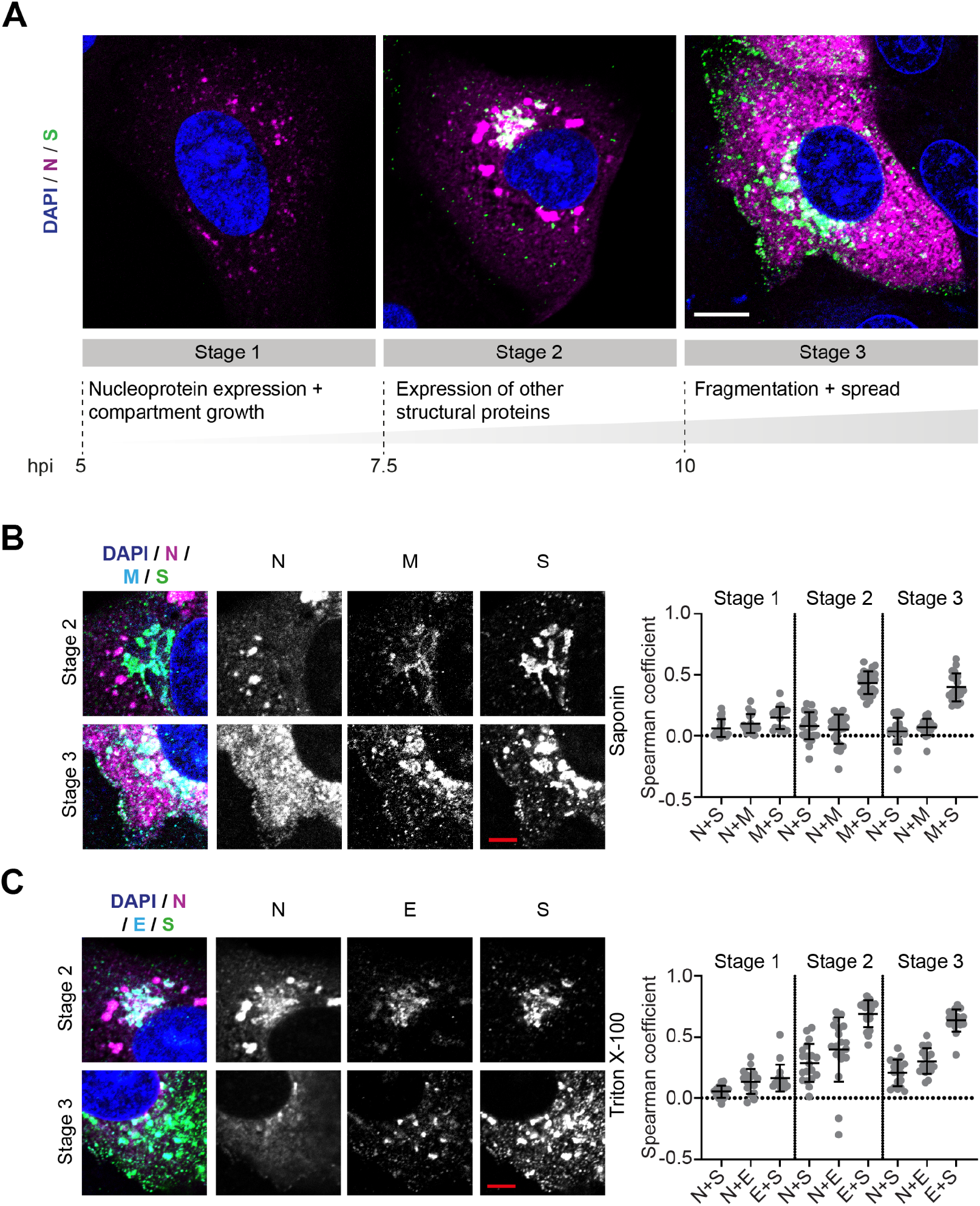
The cellular distribution of structural proteins of SARS-CoV-2 is tightly regulated in space and in time. A) Three categories describing different cell states were identified based on the distribution pattern of the nucleocapsid (N) and spike (S) proteins during the late phase of the SARS-CoV-2 replication cycle. These categories were termed stage 1, 2, and 3, respectively. For each stage, a representative confocal microscopy image is shown. Blue: DAPI-stained nuclei; magenta: nucleocapsid protein (Atto 647N); green: spike protein (Alexa Fluor 488, S). Scale bar 10 μm. B) colocalisation between SARS-CoV-2 nucleocapsid (N), membrane (M) and spike (S) proteins. Left: Representative confocal images of infected Vero cells in infection stages 2 and 3. Blue: DAPI-stained nuclei; magenta: nucleocapsid protein (N); cyan: membrane protein (M); green: spike protein (S). Scale bar: 5 μm. Right: colocalisation between viral proteins N, S and M at different infection stages determined using the Spearman coefficient method (stage 1: n = 20, stage 2: n = 25, stage 3: n = 20). Manders coefficients are shown in Supporting Figure 3A. C) Left: Representative confocal images of SARS-CoV-2 infected Vero cells in infection stages 2 and 3. Cells were fixed with formaldehyde and permeabilized with triton X-100. Blue: DAPI-stained nuclei; magenta: nucleocapsid protein (N); green: spike protein (S); cyan: S envelope protein (E). Scale bar: 5 μm. Right: colocalisation between viral proteins N, S and E at different infection stages determined using the Spearman coefficient method. (Stage 1: n = 16, stage 2: n = 19, stage 3: n = 18). Manders coefficients are shown in Supporting Figure 3B.

The selection of antibodies against the structural proteins allowed us to image three of the four structural proteins at once. The M and E protein could not be visualised simultaneously because the respective antibodies belonged to the same host species. Instead, we immunostained two separate sets of samples, co-staining either for N, M and S proteins (Figure 1B) or N, E and S proteins (Figure 1C). We expected the M and E proteins to show a similar pattern of accumulation in the same host cell membrane compartment as the S protein, as these are all transmembrane proteins. Representative images and quantitative colocalisation analysis confirmed that M (Figure 1B) and E proteins (Figure 1C) co-occurred foremost with the S protein and not the N protein. Consequently, M and E proteins follow the same accumulation and fragmentation pattern as the S protein.

By comparing the colocalisation values between N and transmembrane proteins (M, S and E proteins) within the two separate samples for stages 2 and 3, we noticed that whereas the Spearman coefficients (see Methods 4.16) were close to zero in the first sample (Figure 1B), they were increased in the second sample (Figure 1C). This is not due to a different localisation pattern of the proteins in the separate samples, but due to a different capability to visualise the viral proteins depending on the permeabilization reagents used for immunostaining. Use of strong (Triton X-100) instead of mild (saponin) detergents was required to visualise the N protein at the juxtanuclear membrane compartment - in addition to the bright N protein puncta - where the transmembrane proteins also localize. The colocalisation of N protein with the three transmembrane proteins at this compartment is in line with the current model for SARS-CoV-2 assembly, where viral nucleocapsids are trafficked to membrane compartments enriched with M, S and E proteins for assembly.

### 2.2 The kinetic profile of SARS-CoV-2 replication

In order to assess SARS-CoV-2 replication kinetics, we determined the fraction of cells expressing each of the four structural proteins and the fraction of cells in which double-stranded RNA (dsRNA) was present (indicating initiation of viral RNA transcription) at each time point (Figure 2A). At 5 hpi, 5-10% of cells were positively stained for dsRNA and N protein, but for none of the other structural proteins. Consistently, we detected released viral transcripts in the cell supernatant by RT-qPCR from 5 hpi onwards (Figure 2B). These data confirm the observations of Cortese et al. in Calu-3 cells where PCR, an infectivity assay and transmission electron microscopy (TEM) were employed. In the latter report, the release of viral RNA and infectious virus was observed in parallel with the appearance of DMVs at 6 hours post-infection under similar experimental conditions (*13*). This suggests that the timing of events during viral replication is comparable between Calu-3 and Vero cells. At 7.5 hpi we observed that the fraction of cells positive for N protein increased by up to ~15%, with about a third of the cells also expressing the other three structural proteins. From 10 hpi on, infected cells were expressing all four structural proteins at similar levels. This is again consistent with a significantly increased infectious titre at 10 hpi (Figure 2C), confirming completion of the replication cycle and the production of new viruses. For most cells, it appeared that M and E proteins were expressed simultaneously with the S protein. However, in a few cells, only fluorescence signal from the M, but not the S, protein was detected (~5% of infected cells, see representative cell in Supporting Figure 4). In contrast, E and S proteins always occurred together. This indicates that the M protein is expressed before S and E proteins. At 24 hpi, we observed a doubling in the number of infected cells compared to 12 hpi. We then tracked the average expression level over time by measuring the average fluorescence intensity per cell (Supporting Figure 5). For all four viral proteins the trend was similar: the average expression levels per cell increased until 12 hpi when they saturated. While the average values at 12 hpi and at 24 hpi are comparable, we note that the distributions of values are more homogenous at 24 hpi than they are at 12 hpi.

**Figure 2:**
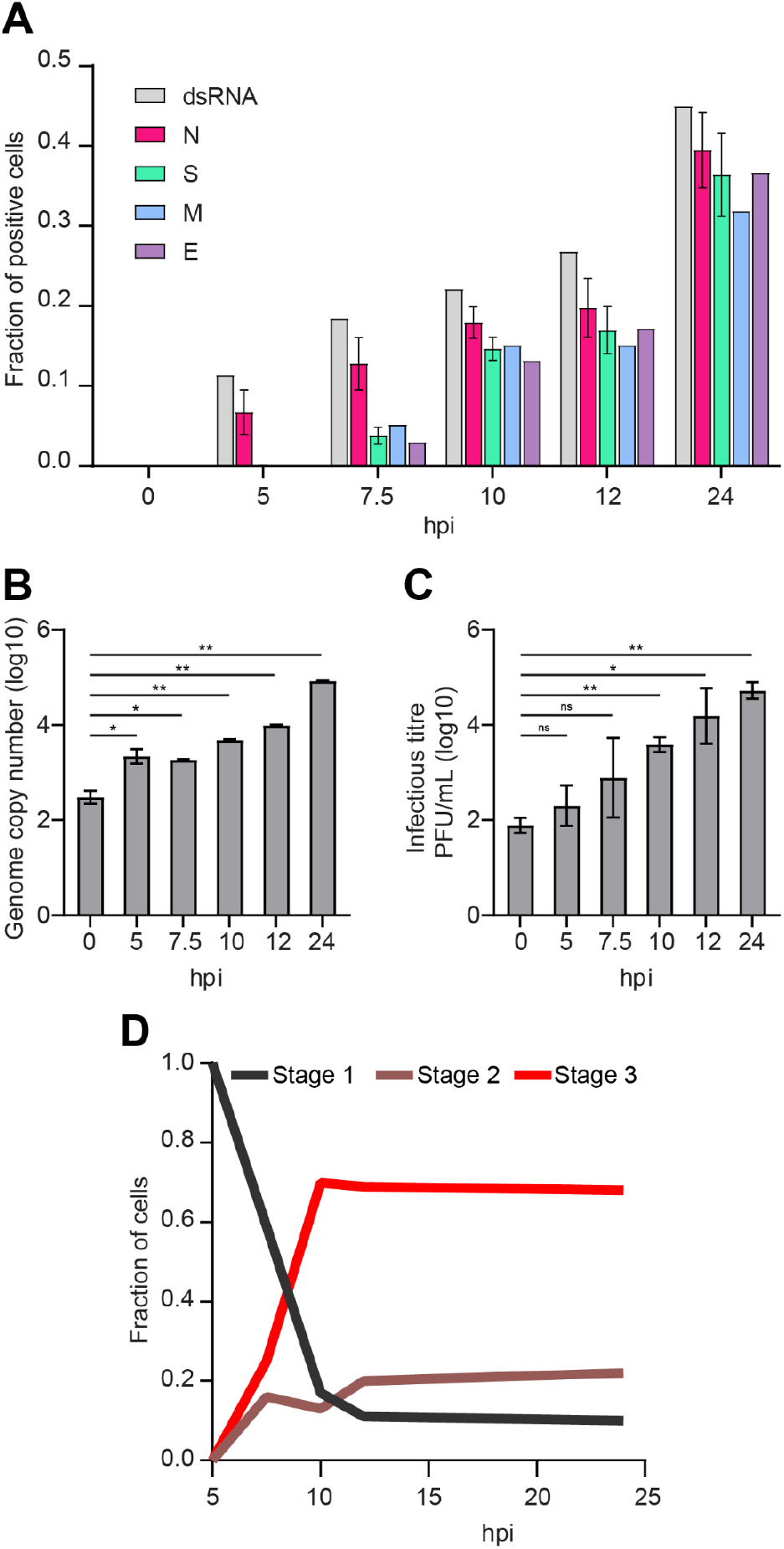
Stepwise expression of the four structural SARS-CoV-2 proteins and dsRNA correlates with staged release of viral transcripts (5 hpi) and infectious virus (7.5 - 10 hpi), respectively. A) The fraction of cells positive for each of the structural proteins of SARS-CoV-2 or viral double-stranded RNA (dsRNA) was determined by immunostaining of infected Vero cells. For each time point, 30-35 widefield microscope images, corresponding to 1000-1500 cells per sample, were analysed. For the control time point of 0 hpi, only around 250 cells were analysed per sample. For the count of N and S positive cells, 3 samples per time point were analysed. B) The copy number of the viral transcripts in the cell supernatant was measured by RT-qPCR. Release of viral RNA was observed from 5 hpi onwards when first cells started expressing nucleocapsid protein. C) The infectious titre of the cell supernatant was determined by the plaque assay. Newly formed infective SARS-CoV-2 virions were released from cells from 10 hours post infection (hpi) onwards. For both assays, two replicates were carried out. Significance was tested with an unpaired t-test. D) Stage-dependent replication kinetics.

Finally, we classified the cells, according to them being in three different stages, to quantify the kinetic profile of the infection process (Figure 2D). Between 5 and 10 hpi, we saw a strong shift from stage 1 to stages 2 and 3. At 7.5 hpi, 50% of cells express all four structural proteins, with an equal number of cells observed in stages 2 and 3 (compact vs. fragmented juxtanuclear membrane compartment). From 10 hpi on, we observe a rapid increase in cells with fragmented compartments (stage 3) which are dominating the population of infected cells (~75%) whereas the fractions of cells in stages 1 and 2 remain low. This indicates that the transition from stage 2 to stage 3 (compact to fragmented juxtanuclear membrane compartment) mainly occurs between 7.5 and 10 hpi. This transition also coincides with a significantly increased production of mature virions at 10 hpi (Figure 2C).

### 2.3 The N protein accumulates around folded ER membranes in convoluted layers that connect to viral RNA replication foci

As shown in Figure 1, the intracellular location of the N protein is distinct from that of the other three structural proteins: initially, the N protein accumulates exclusively in small puncta; as the infection progresses, cytosolic signal gradually increases alongside the puncta (Figure 3A). In parallel, the number, as well as the size, of N protein puncta grow significantly (Supporting Figure 6). At closer inspection of the larger puncta in images of infected cells fixed at 10 hpi, the round structures were found to be shaped like vesicles with an outer layer containing N protein and a hollow center (Figure 3B, left image). We propose that these N protein layers are formed at the viral replication organelles (vRO).

**Figure 3:**
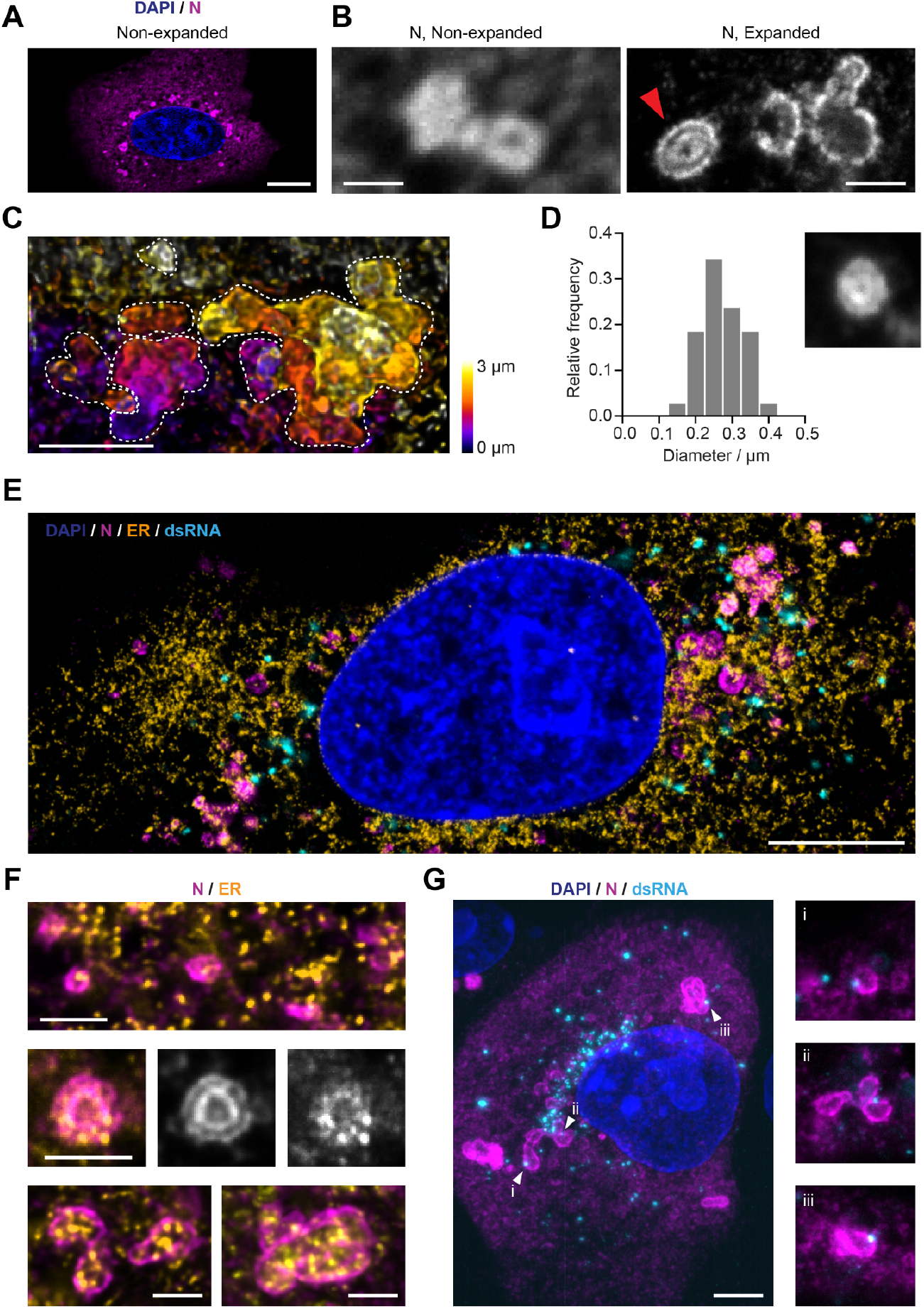
The SARS-CoV-2 nucleocapsid (N) protein is organised in layered structures that host RNA replication foci and are strongly interwoven with the topology of the endoplasmic reticulum. A) Confocal microscopy shows that the nucleocapsid protein forms punctate patterns in the cytosol of infected Vero cells (Blue: nuclei; magenta: nucleocapsid protein). Scale bar 10 μm. The numbers of puncta per cell and size increase in a stage-dependent manner (Supporting Figure 6). B) A combination of expansion and confocal microscopy shows that some of the larger nucleocapsid protein structures observed at later time points (from 10 hpi on) consist of double layers (red arrow). Scale bars 1 μm (taking into account a linear expansion factor of 4.2). C) A combination of expansion and light sheet microscopy reveals convoluted nucleocapsid protein structures, here shown as maximum intensity projections. Each dotted line outlines the boundaries of each convoluted nucleocapsid protein structure. Scale bar 2 μm (taking into account an expansion factor of 4.2). D) Size distribution analysis of the nucleocapsid protein double-layer compartments recorded 12 hours post-infection. The inner circular layer has an average diameter of 275 nm. 38 compartments from 10 cells were analysed. Size of the micrograph is 1.25 μm per side. E) Representative image of an expanded Vero cell stained for the nucleus (blue), ER (calnexin protein, orange), nucleocapsid protein (magenta) and dsRNA (cyan), imaged on a confocal microscope. Scale bar 5 μm (taking into account an expansion factor of 4.2). F) Details of the nucleocapsid protein (magenta) and the ER (yellow), showing that the nucleocapsid protein forms layers around ER membranes. This indicates that the nucleocapsid protein structures might be localised at double membrane vesicles (DMVs) or packets of DMVs. Scale bars 1 μm (taking into account an expansion factor of 4.2). G) A combination of expansion and light sheet microscopy reveals the interaction between dsRNA (cyan) and the nucleocapsid protein N (magenta). This shows that dsRNA foci sit in the layers of the N compartments. Scale bar 5 μm (taking into account an expansion factor of 4.2). Size of the smaller micrographs is 5 μm per side.

To find support for this hypothesis, we applied expansion microscopy (*21*) to investigate these structures in better detail. This super-resolution technique provides a fourfold increase in resolution via a 64x volumetric expansion of the sample. In these higher resolved images of the N puncta we detected that several of the N compartments consisted of double layers of the protein (Figure 3B, right image, and Supporting Video 1). In addition, we observed that single small compartments were often fused to larger convoluted three-dimensional structures (Figure 3C and Supporting Videos 2 and 3).

The inner N protein compartments measured ~275 nm on average in diameter (Figure 3D). Viral replication organelles contain single double-membrane vesicles (DMVs) and DMV packets (VPs) (*15*). DMVs formed by SARS-CoV-2 contained in the ROs are about 300 nm in diameter (*13, 15*), which is in agreement with the structures presented. It is accepted that the DMVs formed by coronaviruses are used by the virus as a protective environment for replication of its RNA genome (*14, 15*), and the presence of SARS-CoV-2 RNA in such structures was recently verified by electron microscopy (*15*).

It has been established that ROs are derived from ER membranes and serve as an anchor for the viral replication and transcription complex (RTC) (*22*). dsRNA is considered a viral replication intermediate indicating the proximity of RTCs. By co-staining the N protein, ER and dsRNA, and then acquiring confocal images of non-expanded cells (Supporting Figures 7A and 7B) as well as expanded cells (Figure 3E), we found that at all stages of infection, the N protein-containing compartments were always associated with the ER. Moreover, the ER membranes seemed clustered at the spots where those compartments are present. In the expanded samples, we observed that the N protein formed layers around the highly convoluted ER membranes (Figure 3F). This was observed for single small (< 1 μm), larger fused (> 1 μm), as well as double-layered N protein compartments.

Analogously to the N protein-enriched compartments, the dsRNA foci were also always associated with the ER (Figure 3E). Only some of the N protein-enriched compartments seemed to colocalise with dsRNA foci whereas many replication foci were not co-occurring with the N protein compartments. A quantitative analysis of confocal images of non-expanded cells showed that the fraction of closely associated compartments and foci decreased for cells in later infection stages (Supporting Figure 8) when the number of dsRNA foci increased (Supporting Figure 7B).

However, single-image cell sections might be misleading, as they omit information either side of the focal plane. In order to analyse the connection between dsRNA foci and N protein-layered compartments, we acquired volume sections of 13 expanded cells using light sheet microscopy (Supporting Video 4). In the majority of samples, most dsRNA foci are located in a region immediately adjacent to the nucleus. We further noted that most N protein compartments were connected to at least one RNA replication focus which was usually situated in the outer layer of the compartment (Figure 3G).

### 2.4 The S protein accumulates in Golgi/ERGIC compartments and transport vesicles containing the lysosome marker LAMP1

Next, we aimed to determine with which organelles the SARS-CoV-2 transmembrane proteins directly interact during assembly and egress. Due to the selection and limitation of the used antibodies, we could only visualise S protein simultaneously with the host cell structures. However, the three transmembrane proteins S, M and E show a high degree of colocalisation making it likely that they behave in a similar fashion.

The S protein was seen to be at least partially located in the Golgi apparatus and ERGIC from a co-occurrence with the respective organelle markers GM130 and LMAN-1 during stages 2 and 3 (Figures 4A and B). This finding corresponds with observations made previously also for SARS-CoV-1 (*23*). We quantified this colocalisation by determining the Spearman coefficients, which exhibited average values of ~0 in control cells and in cells at stage 1, but increased significantly at stages 2 and 3 in all cases.

**Figure 4:**
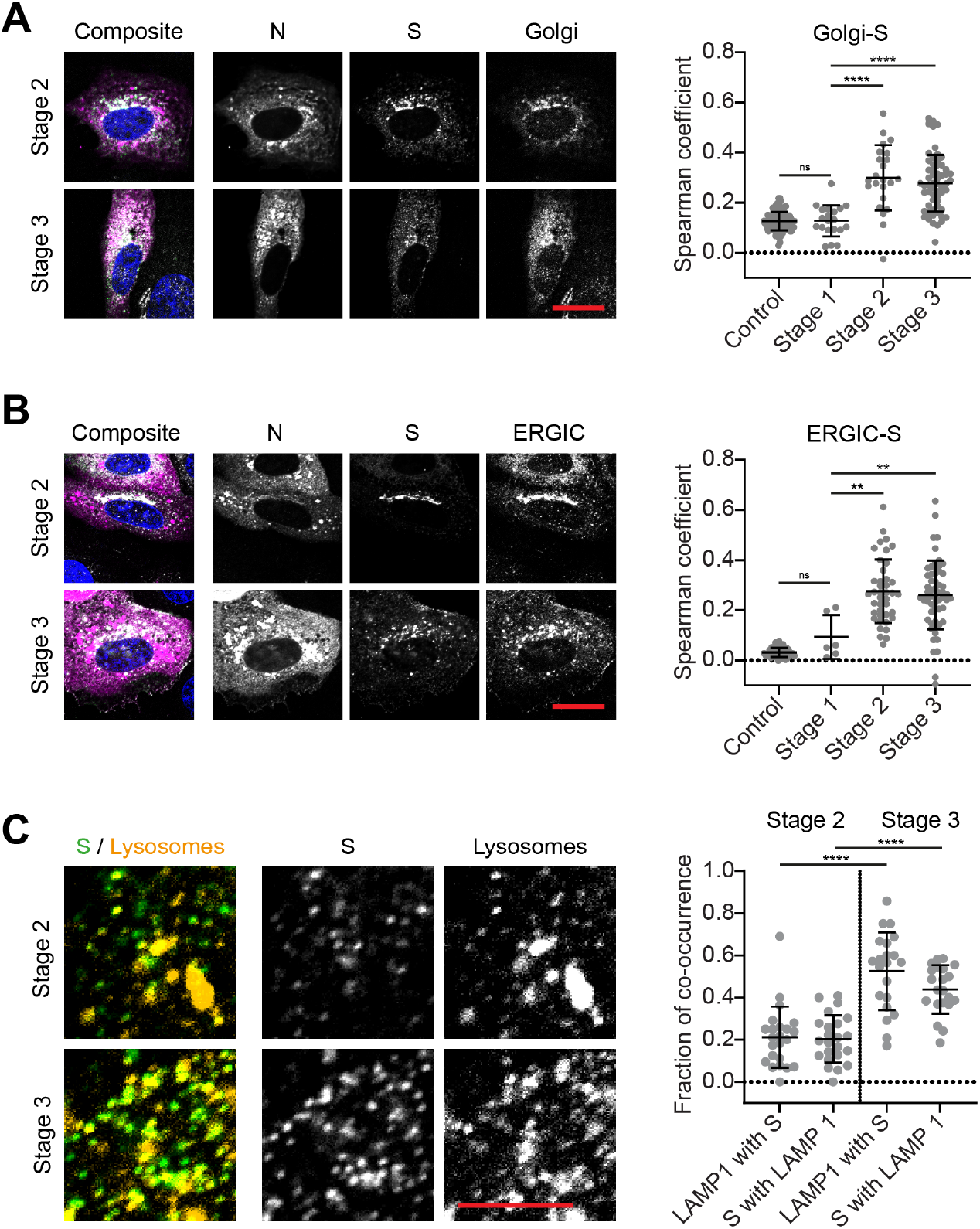
The spike protein (S) accumulates at the Golgi apparatus as well as the ER to Golgi compartment (ERGIC) and colocalises with lysosomes during late infection stages. A) Left: Representative confocal images of SARS-CoV-2 infected Vero cells in infection stage 2 and 3. Blue: DAPI-stained nuclei; magenta: SARS-CoV-2 nucleocapsid protein (N); green: SARS-CoV-2 spike protein (S); cyan: Golgi apparatus (GM130). Scale bar: 20 μm. Right: colocalisation analysis (determined using the Spearman coefficient method) between the Golgi apparatus and the spike protein (S) shows partial spatial correlation from the moment the spike protein starts to be expressed (stage 2) onwards. (Control: n = 111, stage 1: n = 20, stage 2: n = 21, stage 3: n = 59) B) Left: Representative confocal images of SARS-CoV-2 infected Vero cells in infection stage 2 and 3. Blue: DAPI-stained nuclei; magenta: SARS-CoV-2 nucleocapsid protein (N); green: SARS-CoV-2 spike protein (S); cyan: ERGIC (LMAN-1). Scale bar: 20 μm. Right: colocalisation analysis (determined using the Spearman coefficient method) between the ERGIC and the spike protein (S) shows partial spatial correlation from the moment the spike protein starts to be expressed (stage 2) onwards. (Control: n = 44, stage 1: n = 6, stage 2: n = 41, stage 3: n = 58) C) Left: Representative confocal images of SARS-CoV-2 infected Vero cells stained for the spike protein (S, green) and the lysosomes (LAMP1, yellow). Scale bar: 5 μm. Right: Spot-to-spot distance analysis was applied to estimate the fraction of co-occurring spike protein and lysosome spots at stages 2 and 3. Significance was tested for all datasets with an unpaired t-test with Welch’s correction for unequal standard deviations. Manders coefficients are shown in Supporting Figure 9.

Concurrent to the enrichment of S protein at Golgi and ERGIC membranes at stage 2, small spots of S protein appeared in the cytoplasm (Figure 1A and Figure 4A). This indicates trafficking of the S protein, and supposedly also M and E proteins, away from the Golgi and ERGIC membranes. These diffraction-limited spots could either be transport vesicles containing viral proteins in their lipid membranes or newly formed virions. It has recently been reported that lysosomes are used by the virus to exit the cell and that mature virions exploit this for transport to the cell surface (*18*). When we stained the cells for the lysosome marker LAMP1, we found that the spots containing S protein were often also positive for LAMP1 (Figure 4C). We quantified co-occurrence of S protein and LAMP1 for stages 2 and 3 using a spot-to-spot distance analysis. When the centres of spots in both channels were within a distance of 280 nm, they were considered as co-occurring. Interestingly, the fraction of co-occurring spots increased from ~20% to ~50% for both S protein and lysosomes at stage 3. These findings confirm that lysosomes can be used for the shuttling of virions, further supporting the role of lysosomes in SARS-CoV-2 egress.

### 2.5 The infection alters the morphology and location of host cell organelles and cytoskeleton

We further analysed the morphological changes of the host cell organelles involved in SARS-CoV-2 assembly and egress as well as the cytoskeleton at different stages of infection. The most striking morphological change that we noted was a fragmentation of the Golgi compartment. In order to quantify this fragmentation, we measured the angle spanned by the Golgi apparatus around the nucleus, as depicted in Figure 5A. In cells with fragmented compartments, the angles were typically larger than 180°, and often close to 360°. Thus, we distinguished between cells with a compact (<180°) or fragmented (>180°) Golgi compartment. The histograms represent the distributions of angles measured in the cell population at different times post infection. At 5 hpi, we observed almost no fragmentation. At 12 and 24 hpi, however, the fraction of cells with fragmented Golgi compartments was increased. This corresponds to cells in late infection stages (stage 3), when new mature viruses were being produced and released.

**Figure 5:**
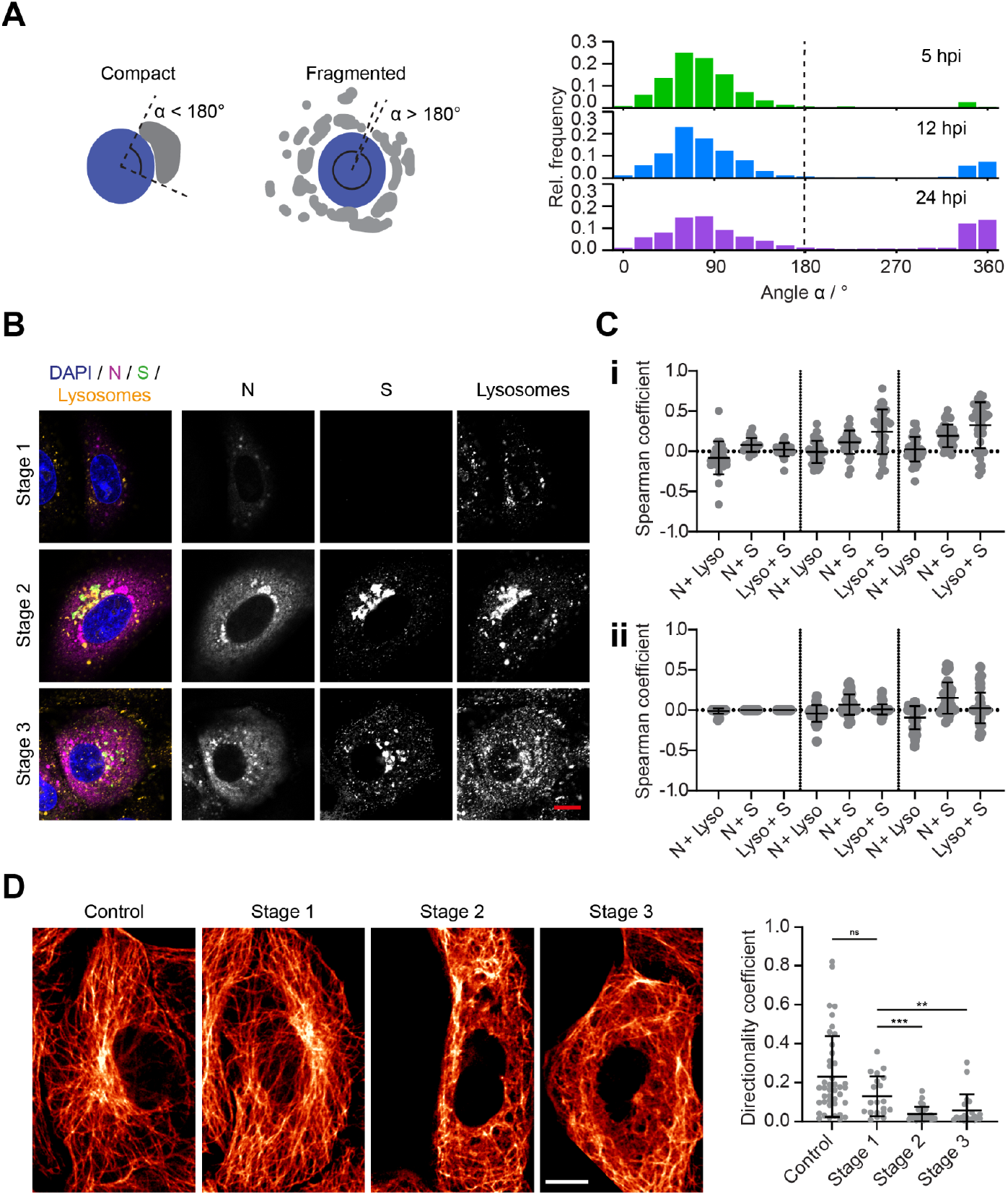
SARS-CoV-2 infection alters the morphology and location of organelles and the cytoskeleton in Vero cells. A) The angular distribution of Golgi compartments around the nucleus was used to distinguish between cells with compact (angle < 180°) and fragmented Golgi apparatus (angle > 180°). The fraction of cells with fragmented Golgi apparatus increased over the time course of infection. The fragmentation of the Golgi apparatus upon SARS-CoV-2 infection was used to sort cells into infection stage 2 or 3 (Figure 2D). B) Representative confocal images of lysosomes in infected Vero cells at different infection stages. Blue: DAPI-stained nuclei; magenta: nucleocapsid protein (N); green: spike protein (S); yellow: lysosomes (LAMP-1 staining). Scale bar: 10 μm. C) colocalisation of the lysosome marker LAMP1 with the nucleocapsid and spike proteins. Otsu thresholding (i) allows correlation between all LAMP1 and SARS-CoV-2 protein signal, whereas manual thresholding (ii) filters out the weaker LAMP1 signal such that only bright, small lysosomal compartments are taken into account (Stage 1: n = 25, stage 2: n = 34, stage 3: n = 39). D) Top: Representative confocal images of microtubules in control (0 hpi) and infected Vero cells (stages 1, 2 and 3). Scale bar 10 μm. Bottom: The directionality coefficient was calculated for subareas of the microtubule network. Each data point corresponds to one subregion inside a cell. Changes in the network due to SARS-CoV-2 infection lead to significant reduction in directionality at stage 1, which is even more pronounced at stage 2 and 3 when all four structural proteins are expressed. Significance was tested with a Mann-Whitney test.

We also noticed that the lysosomes undergo a spatial redistribution during SARS-CoV-2 infection (Figure 5B). At stage 1, the lysosomes were larger on average than in control cells (control: mean area = 0.94 μm^2^, n = 56; stage 1: mean area = 1.22 μm^2^, n = 21), but were still homogeneously distributed within the cytoplasm. When cells started to express S, M and E proteins (stage 2), the lysosome marker LAMP1 was recruited to the Golgi/ERGIC compartments containing the three viral transmembrane proteins, likely through fusion of the lysosomes with the Golgi and ERGIC membranes. The fragmentation of the Golgi apparatus (stage 3) corresponded to a spread of membrane fragments enriched with viral proteins and LAMP1 in the cytoplasm. The correlation of the lysosome signal with the viral proteins N and S (measured via the Spearman coefficient after Otsu thresholding) was moderate at stages 2 and 3 (Figure 5C). There are two distinct sources from which the lysosome signal originates: (i) small and bright compartments resembling the typical lysosome shape, and (ii) LAMP1 accumulated at the Golgi and ERGIC membranes, however with markedly lower intensity. We used manual thresholding to filter out the weaker signal and only investigate the correlation between the small, bright lysosomal compartments with the viral proteins. Interestingly, we detected no correlation between S protein and the lysosomes in that case. Furthermore, there was a negative correlation between N and the lysosomes at stage 3. At this stage, the N protein was widely distributed in the cytoplasm, but excluded from the location of the lysosomes. This indicates that correlation between the viral proteins and LAMP1 only occurs at the compartments after a mixing of membranes.

SARS-CoV-2 infection leads to a remodelling of the microtubule network. Through a directionality analysis, we found that from stage 2 onwards the network loses its orientation (Figure 5D). In non-infected cells, microtubules spread from the microtubule organising center (MTOC) close to the Golgi apparatus to the extremities of the cell. In late infection stages the microtubule filaments were absent from the juxtanuclear area and they were entangled when compared to control cells. For cells in stages 2 and 3, we also detected a loss of cell stiffness which we measured using atomic force microscopy (AFM, Supporting Figure 10). This could be driven by a remodelling of the actin network, which is regarded as the overriding, although not the sole, determinant of cell stiffness (*24*).

## 3 Discussion

We applied advanced fluorescence microscopy to investigate the expression kinetics and spatial arrangement of the four structural SARS-CoV-2 proteins, and studied their interactions with host cell compartments in detail. We observed that the expression of the structural proteins of the virus is tightly staged, with striking differences between N and the three transmembrane proteins. The N protein accumulates mainly in small foci that grow in size and number during the course of infection. Sample expansion in combination with light sheet microscopy revealed that single N protein compartments comprise layered structures of N protein. The compartments resemble complex and convoluted three-dimensional structures as might result from the fusion and engulfment of smaller vesicular subunits. We believe them to be part of the replication organelles formed by SARS-CoV-2. There are several indicators to support this notion. First, the shapes of the N protein compartments resemble those of replication organelles investigated by electron microscopy where interconnected DMVs and vesicle packets were observed (*15*). Second, the smallest structural units we could identify within these convoluted structures were vesicles whose average size was ~275 nm, which matches the size reported for DMVs in Vero cells (*15*). Third, it is known that coronaviruses remodel the host cell ER membranes to integrate the viral replication organelles (*25*). Indeed, we discovered that the N protein-containing compartments are tethered to ER membranes. Finally, we reasoned that if the N protein was associated to the virus replication organelles, the N compartments would be associated with the viral replication and transcription complexes (RTCs). We detected the RTCs by staining of dsRNA, an intermediate of viral RNA replication. Through volumetric imaging, we confirmed that at least one dsRNA focus is usually associated with the outer layer of an N protein compartment, which might consist of several fused sections. We also noted that, similarly to the N protein compartments, the dsRNA foci are always connected to the ER network.

Since one of the functions of the N protein is the encapsulation of the viral RNA, its presence at/around the DMVs and colocalisation with proteins forming the RTCs would not be surprising in accordance with previous reports for SARS-CoV (*16*). However, to our knowledge, association of N protein to ROs has not been reported before, but makes sense to facilitate the spatial organisation of replication and nucleocapsid formation.

If we assume that the N protein-containing structures are indeed part of the virus replication organelles, the question remains of where exactly the N protein is located within those compartments and how this association is formed. It is possible that the protein accumulates in the intermembrane space of the DMV envelope. Another possibility is the accumulation of the N protein at ER membranes while or after they are reshaped into replication organelles. Nonstructural proteins of coronaviruses are known to reshape host cell membranes to induce formation of DMVs (*14*). Either a specific interaction with one or more SARS-CoV-2 nonstructural proteins and/or a curvature-driven binding mechanism could drive an accumulation of N protein. In both cases, accumulation might be affected by a propensity of N protein to phase-separate with RNA (*26–30*). It has been proposed that N protein plays a dual role: the unmodified protein forms a structured oligomer suitable for nucleocapsid assembly, while the phosphorylated protein forms a liquid-like compartment for viral genome processing (*31*). For both processes, association of N protein to the ROs would thus be beneficial.

A study based on cryo-electron tomography showed that strands of naked viral RNA are located within the DMVs (*25*), which are thought to leave the DMVs through a pore in the membrane lining (*17*). However, It is currently not known where and when the newly synthesized viral RNA is encapsidated by the N protein. In the light of the data presented here, we speculate that association of the viral RNA and the N protein to form viral ribonucleocapsid protein complexes (vRNPs) occurs at the membrane of the viral replication organelles. This process might occur either before, or in concurrence with, the release of the newly synthesised RNA into the cytosol. In this sense, we interpret the increasing cytosolic signal of the N protein in late infection stages as an accumulation of vRNPs in the cytosol before and during virus assembly.

Using multi-colour imaging and colocalisation analysis, we show that the SARS-CoV-2 S, M and E proteins all localise at the Golgi and ERGIC compartments in agreement with previous reports (*13*), (*8*). Our study showed furthermore that M protein is recruited to this area slightly earlier than S and E proteins, suggesting a predominant role of M protein for controlling the spatial organisation of the transmembrane proteins and initiating the assembly of SARS-CoV-2. For other coronaviruses, it has indeed been shown that interactions between M proteins form a lattice into which the other two transmembrane proteins of the virus are incorporated (*32*), (*33*), (*34*). Moreover, we detected that the N protein partially accumulates at the Golgi region, however only after expression of the other three structural proteins has taken place. It is known that assembly of coronaviruses is dependent on the M and E proteins, and for SARS-CoV also on the N protein (*34*). In particular, the carboxyl tail of the SARS-CoV M protein interacts specifically with the N protein (*35*). Our results suggest that also for SARS-CoV-2 the M protein is responsible for the recruitment of the N protein to the Golgi/ERGIC membranes.

At late stages of infection, we detected an enrichment of the lysosomal protein LAMP1 at the membrane compartments together with the structural transmembrane proteins of SARS-CoV-2. This suggests a mixing of membranes or a shift/modification in the endolysosomal transport pathways. Immediately after, diffraction-limited spots of S protein can be seen in the cytoplasm. These spots often co-occur with the lysosome marker. Our results corroborate the recent finding that lysosomes are used by coronaviruses for their cell egress (*18*). Therefore, although it is not clear whether these spots contain only vesicles enriched with viral transmembrane proteins or mature virions, the co-occurrence of viral proteins and lysosome markers indicates an immediate onset of the egress pathway after expression of the M, S and E proteins. Interestingly, we detected an increase from ~20% to ~50% co-occurrence at transition from a compact (stage 2) to a fragmented Golgi apparatus (stage 3), indicating a surge in viral egress. It is not clear what causes the fragmentation of the Golgi apparatus. It might be caused by an indirect toxic effect due to the accumulation of viral proteins, by merging of lysosomes with Golgi membranes and/or the manipulation of the microtubule network, which plays an important role in shaping Golgi structure and function (*36*). We found support for the latter by measuring a remarkable rearrangement of the microtubule network after expression of the three SARS-CoV-2 transmembrane proteins, but before Golgi fragmentation occurs. However, further work is needed to elucidate which factors contribute to the defect in the organisation of the Golgi compartments.

We envisage that the methods presented in this study could furthermore be used for studying the role of the non-structural proteins of SARS-CoV-2, the kinetics of the viral genome replication as well as the relationship between the viral RNA, the N protein and the viral replication organelles.

## 4 Material and methods

### 4.1 Biosafety

SARS-CoV-2 was conducted at containment level 3. SARS-CoV-2 infected cells were fixed using previously published and validated protocols (*37*). The results of this experiment were reviewed and approved by the biosafety committee of the Department of Chemical Engineering of the University of Cambridge.

### 4.2 Chemicals

Methanol-free formaldehyde was purchased from Thermo Fisher Scientific; the ampoules were used immediately after opening and any leftover formaldehyde discarded. Glyoxal (40% in water) was purchased from Sigma Aldrich; the glyoxal solution was heated and mixed prior to use to solubilise precipitated glyoxal. Saponin, triton and ammonium chloride were purchased from Sigma Aldrich. All chemicals used for sample expansion (glutaraldehyde 50% in water, sodium acrylate, N,N’-methylenbisacrylamide, acrylamide) were purchased from Sigma Aldrich and used as received. Lyophilised proteinase K was purchased from Thermo Fisher Scientific. Atto 590-conjugated phalloidin was purchased from Sigma Aldrich and solubilised in methanol according to the manufacturer’s instructions.

### 4.3 Antibodies

All the antibodies used in this study are reported in the tables below:

**Table 1:** Primary antibodies.

| Antibody | Supplier | Target | Host species | Dilution |
| --- | --- | --- | --- | --- |
| ab273073 | abcam | Spike protein | Human | 1:400 |
| NB100-56569 | Novus Biologicals | Membrane protein | Rabbit | 1:200 |
| NBP2-41061 | Novus Biologicals | Envelope protein | Rabbit | 1:200 |
| MA1-7403 | Thermo Fisher Scientific | Nucleocapsid protein | Mouse IgG2b | 1:20 |
| Ab01299-2.0 | Absolute antibody | dsRNA | Mouse IgG2a | 1:200 |
| ab22649 | abcam | GM130 | Rabbit | 1:50 |
| ab125006 | abcam | LMAN-1 | Rabbit | 1:50 |
| ab24170 | abcam | LAMP-1 | Rabbit | 1:100 |
| ab22595 | abcam | calnexin | Rabbit | 1:200 |
| ab131205 | abcam | beta-tubulin | Mouse IgG1 | 1:200 |

**Table 2:** Secondary antibodies.

| Antibody | Supplier | Target | Conjugate | Host species | Dilution |
| --- | --- | --- | --- | --- | --- |
| A-11013 | Thermo Fisher Scientific | Human IgG | Alexa Fluor 488 | Goat | 1:200 |
| A-11011 | Thermo Fisher Scientific | Rabbit IgG | Alexa Fluor 568 | Goat | 1:200 |
| A-11031 | Thermo Fisher Scientific | Mouse IgG | Alexa Fluor 568 | Goat | 1:100 |
| A-21144 | Thermo Fisher Scientific | Mouse IgG2b | Alexa Fluor 568 | Goat | 1:200 |
| A-21244 | Thermo Fisher Scientific | Rabbit IgG | Alexa Fluor 647 | Goat | 1:100 |
| 40839 | Merck | Rabbit IgG | Atto 647N | Goat | 1:200 |
| 50185 | Merck | Mouse IgG | Atto 647N | Goat | 1:200 |
| 610-156-040 | Rockland | Mouse IgG1 | Atto 647N | Goat | 1:100 |
| 610-156-041 | Rockland | Mouse IgG2a | Atto 647N | Goat | 1:100 |

### 4.4 Cells and Viruses

Vero cells (ATCC CCL-81) were cultured under standard conditions (37°C and 5% CO_2_) in Dulbecco-modified MEM (Sigma Aldrich) supplemented with 10% heat-inactivated foetal bovine serum (Gibco), antibiotics/antimycotics (100 units/mL penicillin, 100 μg/mL streptomycin, 0.025 μg/mL Gibco Amphotericin B, Gibco) and 2 mM L-glutamine (GlutaMAX, Gibco). Cells were cultured in T-75 polystyrene flasks; splitting took place when cultures reached ~80% cell confluency. For all experiments, cells below passage number 20 were used.

The BetaCoV/Australia/VIC01/2020 strain of SARS-CoV-2 was obtained from the Victorian Infectious Diseases Reference Laboratory, Melbourne (*38*), through Public Health England. This virus was pasaged once in Vero cells for stocks used in this study. The virus was titrated in standard 6-well plaque format on Vero cells and one batch of virus was used for all experiments. Virus sequences were verified by deep sequencing.

### 4.5 Transfection of Vero cells

The four structural proteins of SARS-CoV-2 were expressed in Vero cells using a pEVAC vector backbone. The day before transfection, Vero cells were seeded at 30% confluence in 8-well Ibidi μ-slides (catalogue n. 80826) in antibiotic-free medium. Cells were transfected with Lipofectamine 3000 transfection reagent (Thermo Fisher Scientific) using 100 ng of plasmid DNA and 0.3 μL of Lipofectamine reagent per well. Cells were incubated for 48 hours under standard conditions before fixation and immunostaining as detailed below.

### 4.6 Infection of Vero cells

The day before infection, Vero cells were seeded at 60% confluence in 24-well plates equipped with 13 mm round glass coverslips (VWR, cat n. 631-0150). Cells were washed once with PBS before incubation with BetaCoV/Australia/VIC01/2020 diluted in PBS at an MOI=5. Incubation took place at RT on a rocking plate for one hour, whereupon inocula was removed, cells washed twice with PBS and replenished with complete DMEM. Infection was allowed to progress under standard conditions (37°C, 5% CO_2_) for 0, 5, 7.5, 10, 12 and 24 h time periods. Cells were fixed with either formaldehyde (4% methanol-free formaldehyde in 100 mM cacodylate buffer) or glyoxal (4% glyoxal and 10% ethanol in acetate buffer pH 5, as previously reported (*39*)) after the removal of spent media. Fixation was carried out at 37°C for 20 minutes.

### 4.7 Plaque assay

Plaque assays were performed as previously described for SARS-CoV-1, with minor amendments (*40, 41*). The day before infection, Vero cells were seeded at 30% confluence in 6-well plates. These subconfluent monolayers were infected with 10-fold serial dilutions of each sample in duplicate, diluted in serum-free media, for one hour at R.T on a rocking plate. After removal of the inocula and washing with PBS, 3mL of 0.2% agarose in virus growth media was overlaid and the cells were incubated at 37°C for 72 h. At this time the overlay media was removed, cells were washed with PBS and fixed overnight with 10% formalin. Fixed monolayers were stained with toluidine blue and the plaques were counted manually.

### 4.8 PCR

The viral load of the media collected before cell fixation at 0, 5, 7.5, 10, 12 and 24 h time points post-infection was measured and quantified via quantitative real-time transcription PCR (RT-qPCR). Total RNA extraction of the media was performed using Qiamp viral RNA Mini Kit (Qiagen) following manufacturer’s instructions. 5 μl of the RNA extraction final elution was reverse-transcribed to cDNA and amplified according to the manufacturer’s protocol using TaqMan™ Fast Virus 1-Step Master Mix (ThermoFisher Scientific). The primer pair was as follows: F-5’CAGGTATATGCGCTAGTTATCAGAC-3’ and R-5’CCAAGTGACATAGTGTAGGAATG3’. The probe used was as follows: 5’[6FAM]AGACTAATTCTCCTCGGCGGGCACG[TAM]3’ (Sigma Aldrich). Analysis was performed using the Rotor-Gene 6000 Series Software 1.7 (Corbett Life Sciences, Qiagen).

To generate RNA standards for qRT-PCR, a 97 nucleotide fragment of the spike ORF was cloned into the pJET1.2 vector (Invitrogen). Following linearization with HindIII, in vitro RNA transcripts were generated using the T7 Ribomax Express Large Scale RNA Production System (Promega). Transcripts were purified (RNA Clean and Concentrator, Zymo Research) and the integrity confirmed by gel electrophoresis.

### 4.9 Immunostaining of fixed cells

Cells were fixed as detailed in Section 5.6. The choice of fixative was determined by the structures to be immunostained in each sample, as detailed in Supporting Table 1. A summary of the fixations and permeabilization conditions for the micrographs shown in this paper is reported in Supporting Table 2. Fixed cells were incubated with 50mM NH4Cl in PBS for 10 m to quench fixation. Cells were permeabilized with either 0.2% saponin or 0.2% triton (see Supporting Table 2) in PBS for 15 m and then blocked with 10% goat serum (Abcam) in PBS for 30 minutes (adding 0.2% saponin for saponin-permeabilized samples). Cells were incubated with primary and secondary antibodies for 1 h at RT; antibodies were diluted as detailed in Section 5.3 in PBS containing 1% goat serum (adding 0.2% saponin for saponin-permeabilized samples). Samples not meant for expansion microscopy were counterstained with DAPI (abcam, ab228549) diluted 1:1000 in PBS for 15 m at RT and mounted on glass microscope slides (Fisher Scientific, cat n. 1157-2203) using VectaShield Vibrant mounting reagent (2B Scientific).

### 4.10 Expansion microscopy

The fixed immunostained samples were expanded following a published procedure (*42*) and imaged either on a confocal or on a light sheet microscope as previously reported (*43*). Briefly, immunostained cells were incubated with 0.25% glutaraldehyde in PBS for 15 minutes, washed 3 times with PBS and then incubated with monomer solution (1xPBS, 2M NaCl, 2.5% acrylamide, 0.15% N,N’-methylenebisacrylamide, 8.625% sodium acrylate) for ~2 m at RT. Gelation was started inverting coverslips onto a drop of 150 μL gelling solution (monomer solution/10% TEMED/10% APS, mixed in ratio 96:2:2) and left to gelate for 1 h at RT in a humidified environment. Gels were digested in digestion buffer (1x TAE, 0.5% Triton X-100, 20 mM CaCl2) containing ~8 U/mL proteinase K overnight at 37C. Gels were eventually placed in double-distilled water to expand. The expansion factor (4.2) was calculated as previously reported (*43*).

### 4.11 Microscopes

Widefield microscope: Widefield imaging of fixed SARS-CoV-2-infected cells was carried out on a custom-built automated widefield microscope. Frame (IX83, Olympus), stage (Prior), Z drift compensator (IX3-ZDC2, Olympus), 4-wavelength high-power LED light source (LED4D067, Thorlabs), and camera (Zyla sCMOS, Andor) were controlled by Micro-Manager (*44*). Respective filter cubes for DAPI (filter set 49000-ET-DAPI, Chroma), Alexa Fluor 488 (filter set 49002-ET-EGFP, Chroma), Alexa Fluor 568 (filter set 49008-ET-mCherry, Texas Red, Chroma), Alexa Fluor 647 and Atto647N (excitation filter 628/40, dichroic beamsplitter Di02-R635, emission filter 708/75, Semrock) as well as Atto 490LS (filter set 49003-ET-EYFP, Chroma, emission filter replaced by 600LP, Semrock) were used. Images were acquired with an Olympus PlanApoU 60x/1.42 NA oil objective lens at 30-35 random positions for each sample.

Confocal microscopes: The imaging of non-expanded fixed samples was performed on a Zeiss LSM 800 microscope using a Plan-Apochromat 63x/1.4 NA oil objective. The microscope was controlled using the Zen software (version 2.6) and for acquisition of 16-bit images a pinhole size of 1.0 Airy unit (AU) for each channel, a scan speed of 5 (1.47 μs / pixel) and 4x averaging were used. Pixel size was 70.6 nm. Expanded gels were cut to fit in a round glass-bottom dish (Ibidi μ-dish, cat n. 81158) pre-coated with poly-L-lysine and were imaged on a Leica SP5 microscope using an apochromatic 63x/1.2 NA water objective. Images were acquired using a scanning frequency of 10 Hz and a pixel size ranging from 100 to 150 nm. In order to increase the collection of signal from the samples, the pinhole size was opened to 2.0 AU (in contrast to the preset value of 1.0 AU), which corresponds to an optical section of 1.5 μm.

Light sheet microscope: Expanded samples were imaged on a custom-built inverted selective plane illumination microscope (iSPIM). Parts were purchased from Applied Scientific Instrumentation (ASI) including controller (TG8_BASIC), scanner unit (MM-SCAN_1.2), right-angle objective mounting (SPIM-K2), stage (MS-2K-SPIM) with motorized Z support (100 mm travel range, Dual-LS-100-FTP) and filter wheel (FW-1000-8). All components were controlled by Micro-Manager by means of the diSPIM plugin. The setup was equipped with a 0.3 NA excitation objective (10x, 3.5 mm working distance, Nikon) and a higher, 0.9 NA detection objective (W Plan-Apochromat 63x 2.4 mm working distance, Zeiss) to increase spatial resolution and fluorescence signal collection. Lasers (OBIS445-75 LX, OBIS488-150 LS, OBIS561-150 LS and OBIS647-120 LX, Coherent) were fibre-coupled into the scanner unit. An sCMOS camera (ORCA-Flash 4.0, Hamamatsu) was used to capture fluorescence. Respective emission filters were BrightLineFF01-474/27, BrightLineFF01-540/50, BrightLineFF01-609/54 and BrightLineFF0-708/75 (Semrock). Gels containing expanded samples were cut into small strips and mounted onto 24×50 mm rectangular coverslips with expanded cells facing upwards using Loctite super glue (Henkel), as previously reported (*43*). The sample was then placed into an imaging chamber (ASI, I-3078-2450), which was filled with double-distilled water. We recorded volumes with planes spacing 0.5 μm. Raw data were deskewed using a custom MATLAB routine including a denoising step to remove hot pixels. Stacks were automatically separated in the respective colour channels and individually processed. Maximum intensity projections were generated of the deskewed stacks.

Correlative structured illumination and atomic force microscope: Correlative atomic force/fluorescence microscopy measurements were performed as described before (*45*). Atomic force microscopy (AFM) measurements were performed on a Bioscope Resolve AFM (Bruker), operated in PeakForce QNM mode, which was combined with a custom-built structured illumination microscopy system (*46*). A 60x/1.2 NA water immersion lens (UPLSAPO 60XW, Olympus) was used for fluorescence excitation and detection which was captured with an sCMOS camera (C11440, Hamamatsu). The wavelengths used for excitation were 488 nm (iBEAM-SMART-488, Toptica), 561 nm (OBIS 561, Coherent) and 640 (MLD, Cobolt). Images were acquired using customized SIM software.

### 4.12 Deconvolution

Confocal images and deskewed light sheet microscopy data of expanded samples were deconvolved using the PSF Generator and DeconvolutionLab2 plugins in Fiji (*47*). In total, 25-100 iterations of the Richardson–Lucy algorithm were used. Deconvolved data were maximum intensity projected in Fiji, optionally using color to indicate depth.

### 4.13 Replication kinetics from widefield data

The percentage of cells expressing each of the structural proteins of SARS-CoV-2 and dsRNA was calculated semi-automatically using the image processing program Fiji (*48*). Cells expressing SARS-CoV-2 proteins and dsRNA were counted manually whereas the total number of cells was determined automatically using the ‘Analyse particles’ function. Images of the DAPI-stained nuclei were filtered using the ‘Subtract background’ rolling ball radius = 20 pixels), ‘Gaussian blur’ (sigma =15 pixels) and ‘Unsharp Mask’ (radius =10 pixels, mask weight = 0.8) functions. Otsu thresholding was used to create a binary mask image. Dividing cells and cells at the edges of the image were excluded from analysis. For each time point, 30-35 widefield microscope images (1000-1500 cells) were counted. For the control time point 0 hpi, only around 250 cells were counted. In order to determine the average expression levels of each of the structural proteins of SARS-CoV-2, the infected cells were segmented manually and the average fluorescence intensity in each viral protein channel was measured. From each value, the mean background intensity was subtracted and data normalised using the highest average intensity value of the respective time point (usually at 12 or 24 hpi) for each protein. For each time point and protein, more than 80 cells were analysed, except the early time point of 5 hpi with only around 40 cells since the fraction of infected cells was very low. (5 hpi n = 46, 7.5 hpi: n = 84, 10 hpi: n = 85, 12 hpi: n = 88, 24 hpi: n= 81 for N, S and M, and 5 hpi n = 40, 7.5 hpi: n = 104, 10 hpi: n = 83, 12 hpi: n = 94, 24 hpi: n = 82 for E).

### 4.14 Stage kinetics from widefield data

The OpenCV and scikit-image Python libraries were used for analysis. Quantification was performed on a dataset of ~35 widefield images per time point stained for the SARS-CoV-2 nucleocapsid protein, the SARS-CoV-2 spike protein, GM130 (Golgi apparatus) and nucleus (DAPI). Infection stages were assigned to each infected cell in the following way. First, binary masks for the cell nuclei were created by using local Otsu thresholding followed by contour detection and filtering (see Section 4.18). For nucleocapsid protein detection, global Otsu thresholding was applied to the corresponding channel. Then, for each detected nucleus, the nucleus masks were used to create thin perinuclear regions around the edge of each nucleus by upscaling each mask by a factor of 1.15 and subtracting the original mask, producing thin hoops around each nucleus. The cell was counted as containing the nucleocapsid protein if more than one pixel in this region was above the threshold value. The spike protein analysis was the same, except a fixed threshold value was used instead of Otsu thresholding, and the masks were upscaled by a factor of 1.2 to produce a thicker perinuclear region, as spike protein signal was more sparse than that of the nucleocapsid protein. If cells were positive for nucleocapsid protein, but not spike protein, they were classified as stage 1. In order to distinguish between stages 2 and 3, a fragmentation analysis of the Golgi apparatus (see Section 4.18) was performed.

The accuracy of the method was checked by manually counting the fraction of cells with nucleocapsid protein and spike protein signal as well as the fraction with a fragmented Golgi apparatus at ~10-15 images at 5 hpi and 10 hpi. The results produced by the algorithm were within one standard deviation of the mean values determined by manual analysis.

### 4.15 Image segmentation

The OpenCV and scikit-image Python libraries were used for the segmentation. Widefield and confocal images were segmented to enable cell-specific analysis of the dataset as follows. Initially, all channels of the images were merged to a grayscale image and background was removed via Li thresholding (*49*). Connected component analysis was performed to segment the single cell units in the image. To be segmented as an object of interest, a connected cluster was filtered via a minimum size of ~150 μm^2^ (corresponds to 30.000 pixels for confocal images, the size of a typical cell nucleus was ~200-250 μm^2^). In case of high cell density or staining of extended structures (e.g. microtubules), connected component analysis might lead to large numbers of cells being detected as one cluster. Here, when a maximum cell cluster size of 1.000.000 pixels was extended, the number of nuclei in the cluster was isolated using the DAPI-stained nuclei. For each nucleus in the cluster, the distances to its K nearest neighbouring nuclei were measured (usually use K = 2 or K = 3, given that in most cases <10 cells make up one cluster). The cell outline of each single cell unit was then defined by the outlines of the nucleus (DAPI-channel) and the half-distances to its K nearest neighbours (choosing the maximum sized box that included all mentioned positions). For images characterised by low cell density, the described methods successfully segmented all cells that can be identified manually. For high cell density images or including extended cell structures, these methods led to a good estimation of the cell outline for the majority of cells (>75% by visual inspection).

### 4.16 colocalisation analysis

We quantified the spatial correlation between all four viral structural proteins by measuring Spearman’s rank coefficients. The Spearman coefficient is based on the ranking of image intensities. After assigning ranks to the pixel intensity values in each of the two channels, the Pearson correlation, which measures the degree of correlative variation, between the rank values of the pixel intensities in the two images is calculated. We also calculated the Manders coefficient which in contrast to the Spearman coefficient measures co-occurrence of intensities in the two channels rather than their correlation (*50*). However, interpretation of the Manders coefficient can be difficult since it depends on the ratio of total intensities in both channels. In contrast to the Spearman coefficient, the Manders coefficient is also affected by out-of-focus signal.

Spearman’s rank coefficients and Manders overlap coefficients were computed by ColocAnalyzer. ColocAnalyzer is a custom program for image filtering and colocalisation analysis, which is free and available here: https://github.com/LAG-MNG-CambridgeUniversity/ColocAnalyzer. Firstly, we saved images in such a manner that each of the channels of interest fell into one of three main colors: red, green or blue. Then, we chose the two channels of interest (for example red+blue or green+red) to be analysed. For each image, Otsu thresholding was applied before computing colocalisation coefficients on the remaining pixels with higher intensities.

Spearman’s rank coefficients were computed by ColocAnalyzer as:

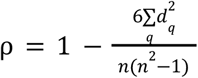

Here *d* = *rank*(*I*1(*q*)) – *rank*(*I*2(*q*)) is the difference between ranks computed for pixel *q* in channel 1 and in channel 2 independently. *n* is the number of pixels that were analysed.

Since after thresholding a significant fraction of pixels was blanked (would have zero intensity), we used only those pixels that had non-zero values in both channels to avoid an impact from black pixels.

The Manders overlap coefficient was computed by ColocAnalyzer using the formula provided in the original paper (*51*):

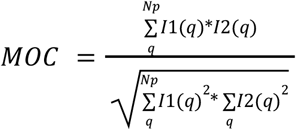

Where *I*1(*q*), *I*2(*q*) are the intensities of pixel*q*in the first and second channel respectively. *Np* is the total number of pixels taken for analysis.

### 4.17 Spot detection and analysis

Nucleocapsid protein, dsRNA and lysosome spot detection and analysis from microscopy images were performed by a customized MatLab routine. For spot detection, we first applied median filtering to the image: each pixel intensity value is decreased by a median value of intensities in a subarea of 60×60 pixels around this pixel (the size of this subarea was chosen empirically). After Otsu thresholding of the filtered image, we determined the positions of connected pixels with non-zero intensities. We called each cluster of such connected pixels a spot. Finally, we filtered out spots which were smaller than 300 nm in diameter (approximately corresponds to Abbe’s resolution limit) and had a mean intensity value smaller than 10% of the maximum intensity. The area of each detected spot was calculated from the number of pixels per spot (pixel size was 107.3 nm). For spot shape analysis, we fitted each spot to an ellipse with the customized MatLab routine “fit_ellipse.m” (https://www.mathworks.com/matlabcentral/fileexchange/3215-fit_ellipse), and used the two radii *R_min_* and *R_max_* obtained from fitting to compute the eccentricity value of each spot: 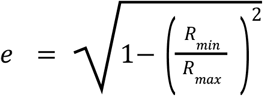, where *R_min_* and *R_max_* are the smaller and larger radii, respectively.

The distance between nucleocapsid protein and dsRNA spots was calculated as the minimal distance between the two spot centers.

Spot-to-spot distance between spike protein and the lysosome marker LAMP1 was analysed using the spot colocalisation plugin ComDet for Fiji. For particle detection within the plugin, particle sizes between 3 and 4 pixels (corresponds to 210-280 nm) and an intensity threshold of 3-10 standard deviations of the average particle intensity were selected. The maximum distance between colocalized spots was set to 4 pixels (corresponding to 280 nm).

### 4.18 Fragmentation analysis

Fragmentation analysis of the Golgi apparatus was performed on 16-bit widefield images. The OpenCV and scikit-image Python libraries were used for the analysis. First, the channel with the DAPI-stained nuclei was segmented into cell nuclei and background using local Otsu thresholding followed by contour detection using the cv2.findContours function within the OpenCV library. The detected contours were filtered by size and circularity to ensure only the single non-overlapping nuclei were selected. Specifically, contours with lengths in the range of 250-2500 pixels (30-300 μm) were accepted. From manual inspection, the nuclei contours fell roughly within the range of 300-700 pixels (35-85 μm). Only contours with length-to-area ratios below 0.05 were selected to eliminate non-elliptical shapes. The contours were then scaled down to 90% of their original size to avoid overlap with structures from other channels and filled to produce a mask for each image.

Next, local Otsu thresholding was performed on the Golgi apparatus channel. The mask of the corresponding nucleus was subtracted from the result. A rectangular region was created around each detected nucleus for subsequent location of the Golgi apparatus. The size of the region was determined by first creating a rectangle such that its borders were tangential to the outline of the detected nucleus and then scaling up its size by a factor of 2. Contour detection was performed within each region to locate the Golgi apparatus or its fragments. The angular size of each contour with respect to the centre of the corresponding nucleus was calculated. It was found that a fragmented Golgi apparatus was typically detected as a single contour since thresholding of the widefield images did not resolve the large number of small fragments, so only the size of the largest detected fragment for each corresponding nucleus was recorded. Contours with the angular size below 20° were found to be indistinguishable from noise, and so the corresponding cells were excluded from the analysis.

### 4.19 Microtubule directionality analysis

Directionality of microtubules was computed by a custom MATLAB routine based on the texture detection technique (TeDT) introduced in (*52*). The method relies on computing gray-level co-occurrence matrices (GLCMs) as proposed in (*53*). The matrix is defined for single values of pixel position shifts [*dx, dy*] and consists of relative frequencies *p_ij_* that two pixels with gray levels *i* and *j* are separated by [*dx, dy*]. For 8-bit images, the GLCM will be a matrix of 256 x 256 elements. Instead of using [*dx, dy*], we used the concept of angle and distance: [*φ, d*]. We varied the distances from 10 pixels to 100 pixels in 5 pixel steps (19 values). The minimum of 10 pixels corresponds to approximately 1 micron, so that short microtubules less than 1 micron in length were excluded from the analysis. The range of directions *φ* = [0°: 180°] was divided into 45 segments with 4°steps for fine resolution of directionality. In total, we generated 19 x 45 = 855 GLCM matrices for each image. Then, as in (45), we computed the joint probability of occurrence for the specified pixel pair:

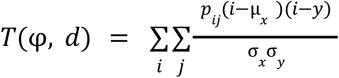

where μ_*x*_, μ_*y*_, σ_*x*_, σ_*y*_ are the means and standard deviations of *p_x_* and 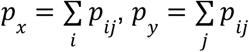.

Next, we averaged those values across distances to leave only the angular dependence:

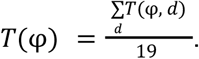

Then we obtained the texture correlation values *H*(*φ*) by normalizing the joint probability for each direction:

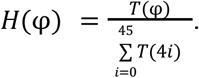

The texture correlation function shows greater values for the angles with preferable directions in microtubule images. Visual inspection on a number of microtubule images showed good performance of the method and its ability to find precisely (up to 4^°^ in our case) dominating microtubule directions in the image. Finally, the directionality coefficient was computed from summing up the second moments around each peak, from valley to valley:

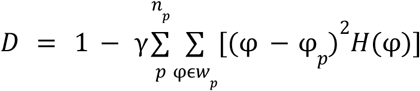

where *n_p_* - number of peaks in *H*(*φ*), *φ_p_* - value of an angle at the *p*-th peak, *w_p_* range for *p*-th peak between two valleys and *γ* - normalizing coefficient: 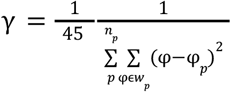.

### 4.20 Cell stiffness measurement and analysis

For AFM cell stiffness measurements, Vero cells were plated at 60% confluence in 50 mm glass bottom dishes (GWST-5040, Willco Wells BV) the day before infection, and infected and fixed as described before. Live Cell probes (PFQNM-LC, Bruker AFM probes) were used for all experiments. The probes were pre-calibrated for spring constant (nominal 0.08 N/m) and deflection sensitivity was calibrated at the start of each experiment. The force applied to the cells was kept constant throughout the experiments, with typical values ranging between 150 - 300 pN. Force curves were fitted to a Hertz model:

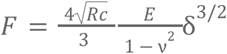

where *R_c_* is the radius of tip curvature, *υ* is the sample’s Poisson’s ratio, *E* is the Young’s Modulus, and *δ* is the indentation depth. Curve fitting and Young’s modulus calculation were performed using Nanoscope analysis.

## Author contributions

C.F.K, K.M.S and L.M conceptualised the project. *K.M.S and L.M contributed equally. G.W.C prepared samples for all experimental data. K.M.S, L.M, L.C.S.W and A.F.-V stained all samples and performed all imaging experiments. G.W.C, H.S, M.S.S and C.G performed plaque assays and PCR experiments. K.M.S, L.M, S.M, M.B and M.B performed data analysis. I.M performed and evaluated AFM experiments. J.R.L provided code for light sheet data processing. K.M.S, L.M, G.W.C, L.C.S.W, S.M, M.B, M.B. I.M, H.S and C.F.K contributed to manuscript writing. All authors revised the article.

## Acknowledgements

We thank Rachel Ulferts for her help in selecting the SARS-CoV-2 antibodies and James D. Manton for his help with the alignment of the light sheet microscope.

## Supporting information

**Supporting Figure 1:**
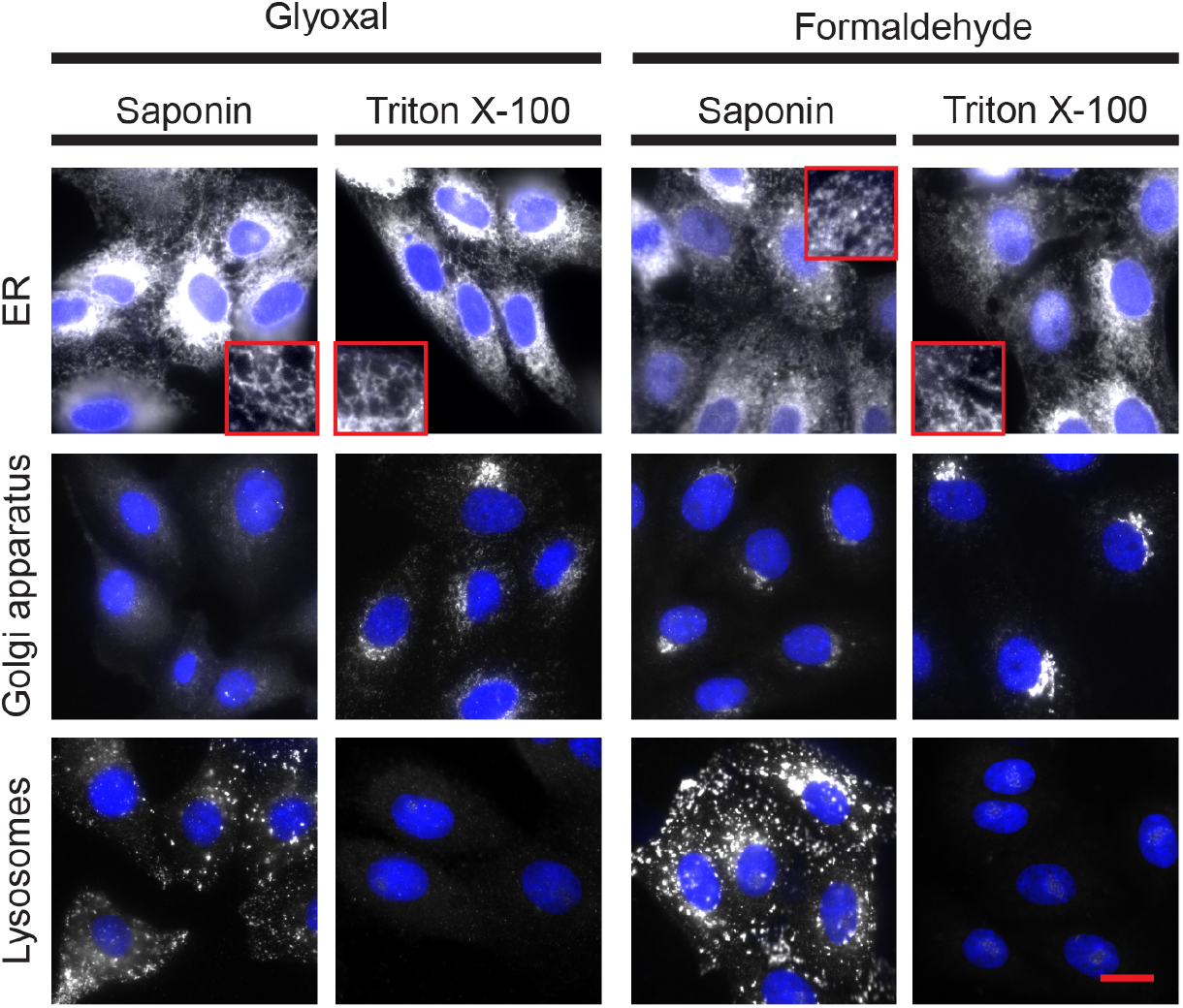
Optimal fixation and immunostaining conditions differ for host cell organelles. The immunostaining of three different cell structures (endoplasmic reticulum (ER), Golgi apparatus and lysosomes) in fixed Vero cells using different chemical fixatives (formaldehyde, glyoxal) and different detergents (triton X-100, saponin) shows that each structure has its own optimal fixation and permeabilization conditions. The ER was stained using an anti-calnexin antibody; the Golgi apparatus was stained using an anti-GM130 antibody; the lysosomes were stained using an anti-LAMP1 antibody. All primary antibodies were detected using the same Alexa Fluor 488-conjugated secondary antibody. White channel: cellular structures; blue channel: cell nuclei. In the red square: detail of the tubular region of the ER. All pictures shown were acquired on a custom-built widefield microscope. Scale bar: 20 μm. The size of the micrograph inserts is 12 μm per side.

**Supporting Figure 2:**
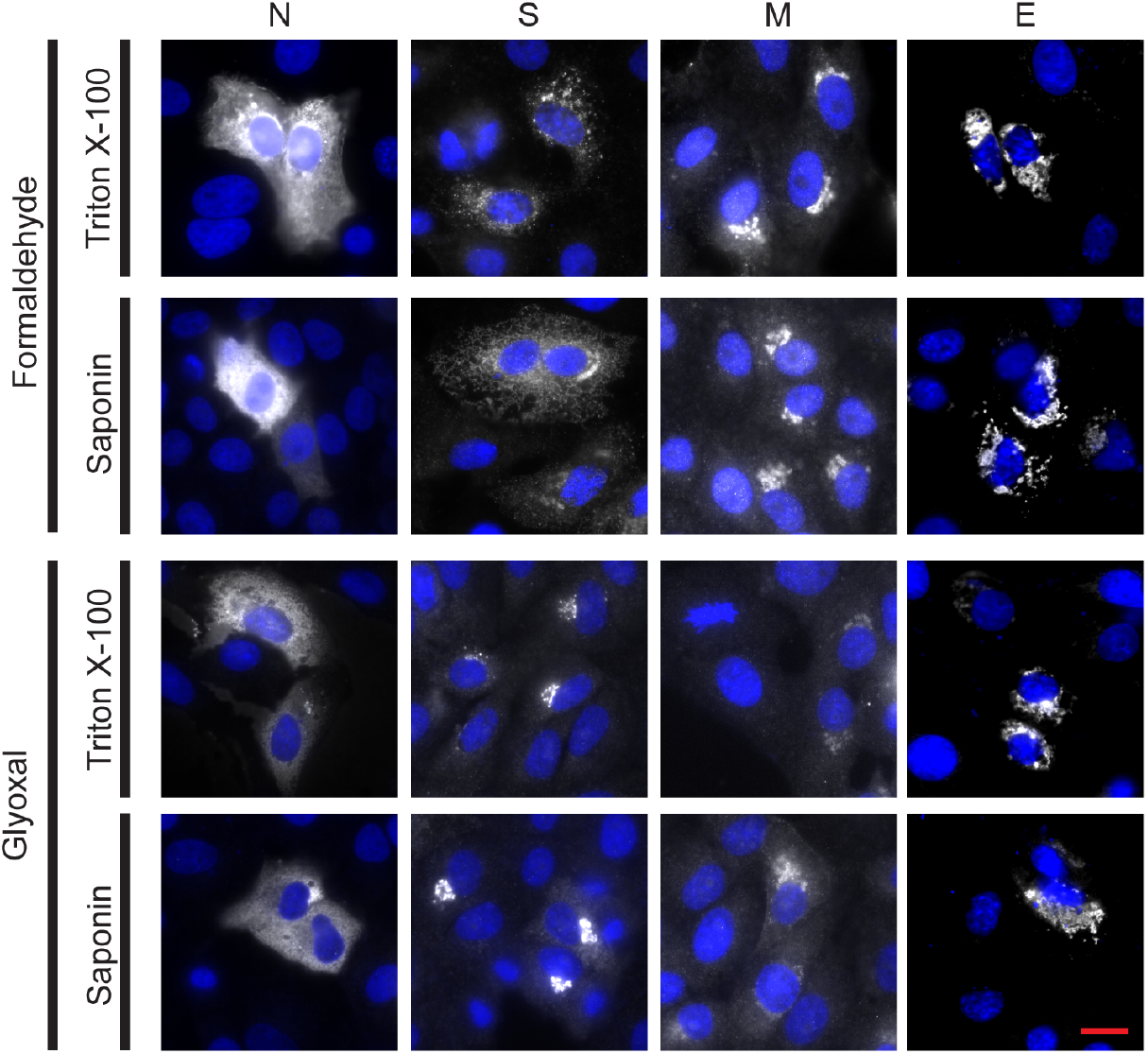
Immunostaining of the four structural proteins of SARS-CoV-2 (nucleocapsid (N), spike (S), membrane (M) and envelope (E)) was successful for all fixation conditions and selected antibodies. Vero cells were transfected to express each of the four viral proteins. The immunostaining was successful for all chemical fixation (formaldehyde or glyoxal) and permeabilization (triton X-100 or saponin) conditions tested. The spike protein pattern was different in the two fixation conditions tested, although this difference was not noticed in the infected samples. Nucleocapsid protein was detected using an Alexa Fluor 568-conjugated secondary antibody; spike, membrane and envelope proteins were detected using Alexa Fluor 488-conjugated secondary antibodies. White channel: SARS-CoV-2 proteins; blue channel: DAPI-stained nuclei. All pictures shown were acquired on a custom-built widefield microscope. Scale bar: 20 μm.

**Supporting Figure 3:**
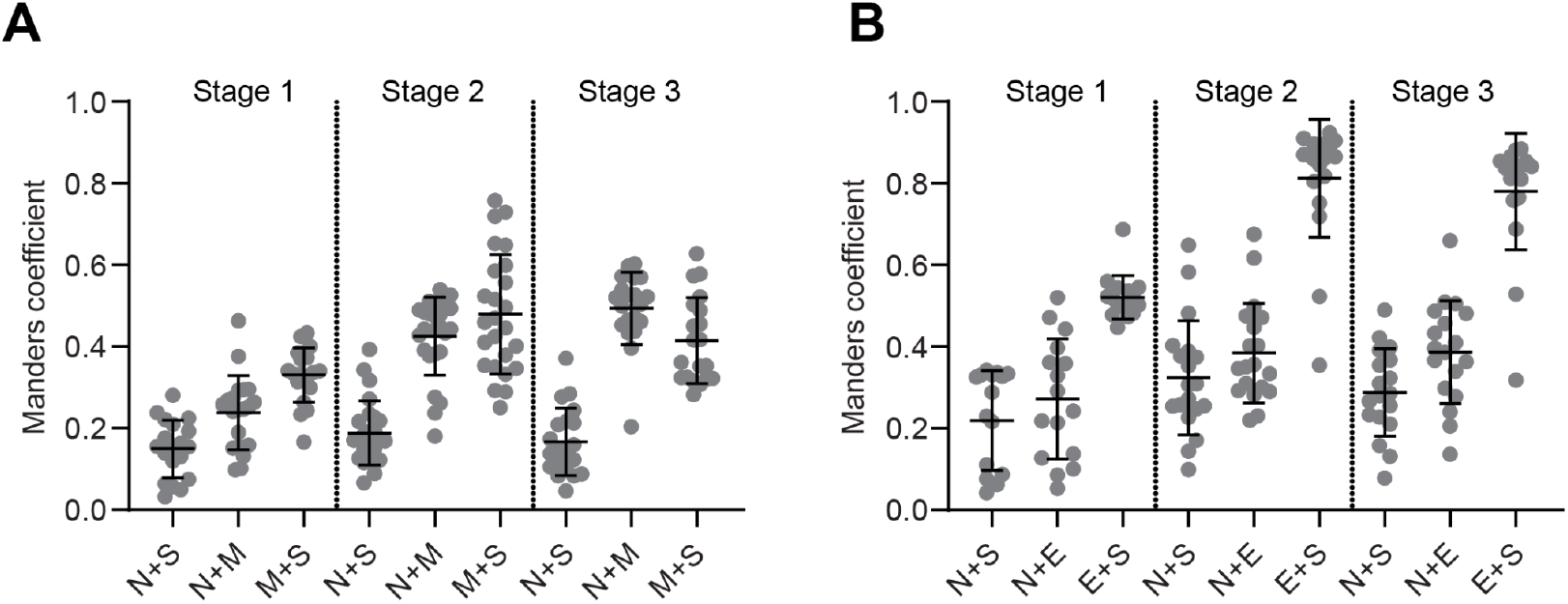
The SARS-CoV-2 spike (S), membrane (M) and envelope (E) proteins highly colocalize, while the nucleocapsid (N) protein shows lower colocalisation with the other viral proteins. A) colocalisation analysis between N, S and M at different infection stages determined using the Manders coefficient method. B) colocalisation analysis between N, S and E at different infection stages determined using the Manders coefficient method.

**Supporting Figure 4:**
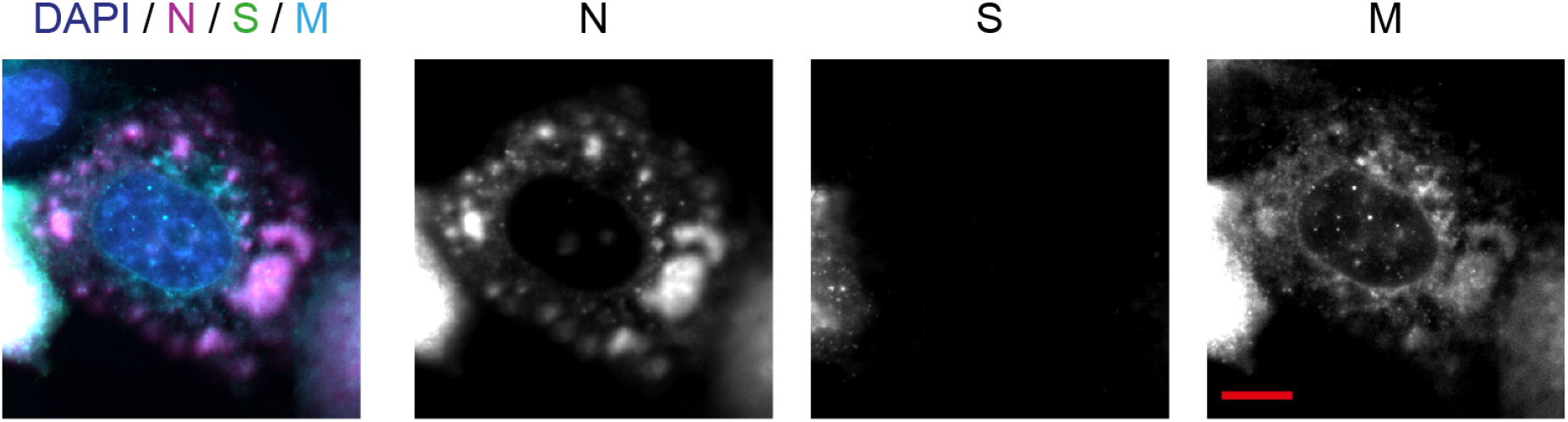
The SARS-Cov-2 membrane protein is expressed slightly earlier than the spike protein in infected Vero cells. In ~5% of cases, infected cells were expressing the membrane protein but not the spike protein, indicating slightly different expression kinetics. Images were acquired on a custom-built widefield microscope. Magenta: nucleocapsid protein (N); green: spike protein (S); cyan: membrane protein (M); blue: nuclei. Scale bar: 10 μm.

**Supporting Figure 5:**
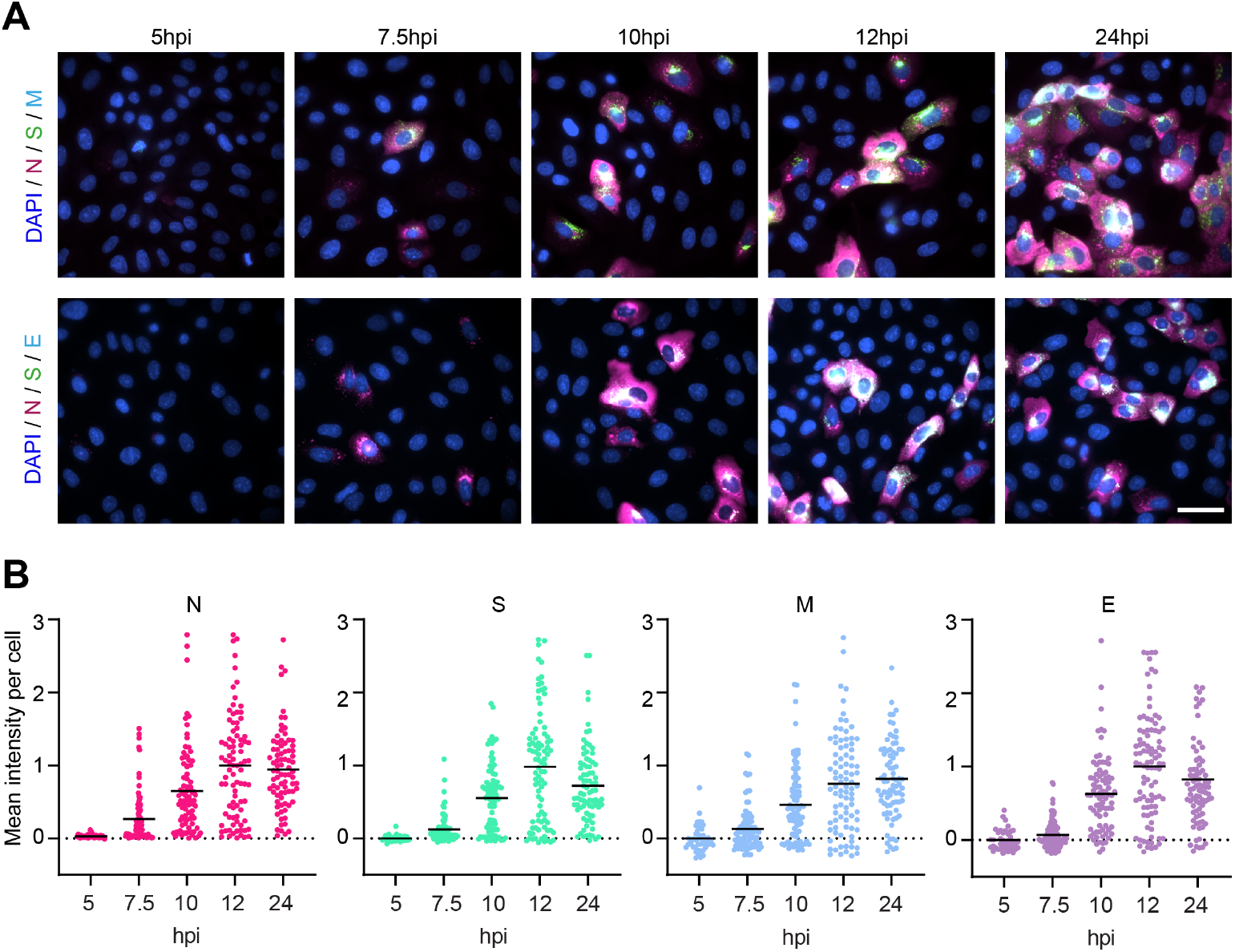
Expression of the four structural SARS-CoV-2 proteins (N, S, M and E) in infected Vero cells saturates at 10 hpi. A) Representative widefield microscopy images of SARS-CoV-2 infected Vero cells, fixed and immunostained at a series of time points post infection. Magenta: nucleocapsid protein (N); green: spike protein (S); cyan: membrane protein (M); blue: nuclei. Scale bar 50 μm. B) The images were used to determine the expression levels of each of the SARS-CoV-2 structural proteins by measuring the average fluorescence intensity per infected cell. Each data point in the graphs represents one cell. N is expressed from 5 hpi onwards, while the other proteins (S, M, E) are expressed concurrently from 7 hpi. The expression of all four structural proteins increases almost linearly up to 12 hpi, when it plateaus, suggesting an equilibrium between newly synthesized viral proteins and released virions.

**Supporting Figure 6:**
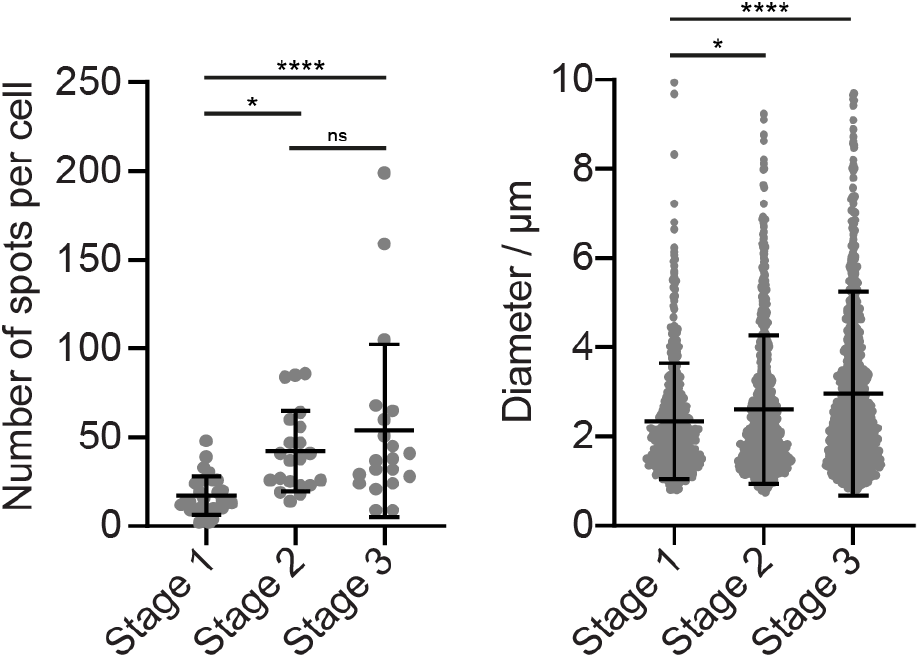
The average number of N compartments per cell as well as their size increase as the infection progresses. Left graph: stage 1: n = 30, stage 2: n = 21, stage 3: n = 30. Right graph: stage 1: n = 515, stage 2: n = 887, stage 3: n = 1076.

**Supporting Figure 7:**
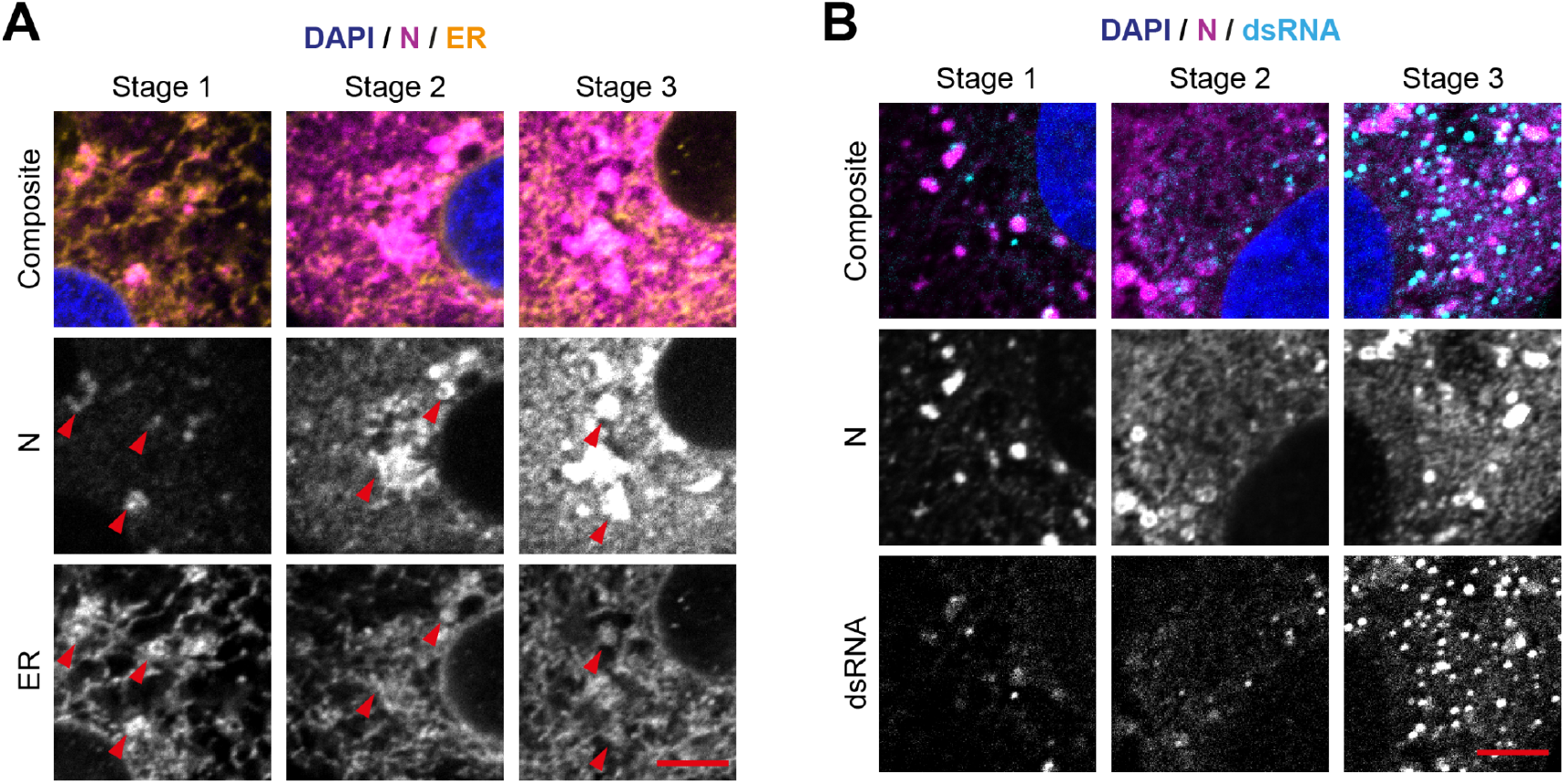
Nucleocapsid protein (N) foci are connected to the endoplasmic reticulum increase, but only partially with RNA replication foci. A) Confocal images of cells in all three infection stages show that the N compartments (magenta) form at regions of condensed ER (orange) and are tethered to it at all stages of infection. Scale bar 5 μm. B) Confocal images of cells in all three infection stages show that the RNA replication foci (cyan) increase in number as the infection progresses and that they partially overlap with the compartments formed by the nucleocapsid protein (magenta). Scale bar 5 μm.

**Supporting Figure 8:**
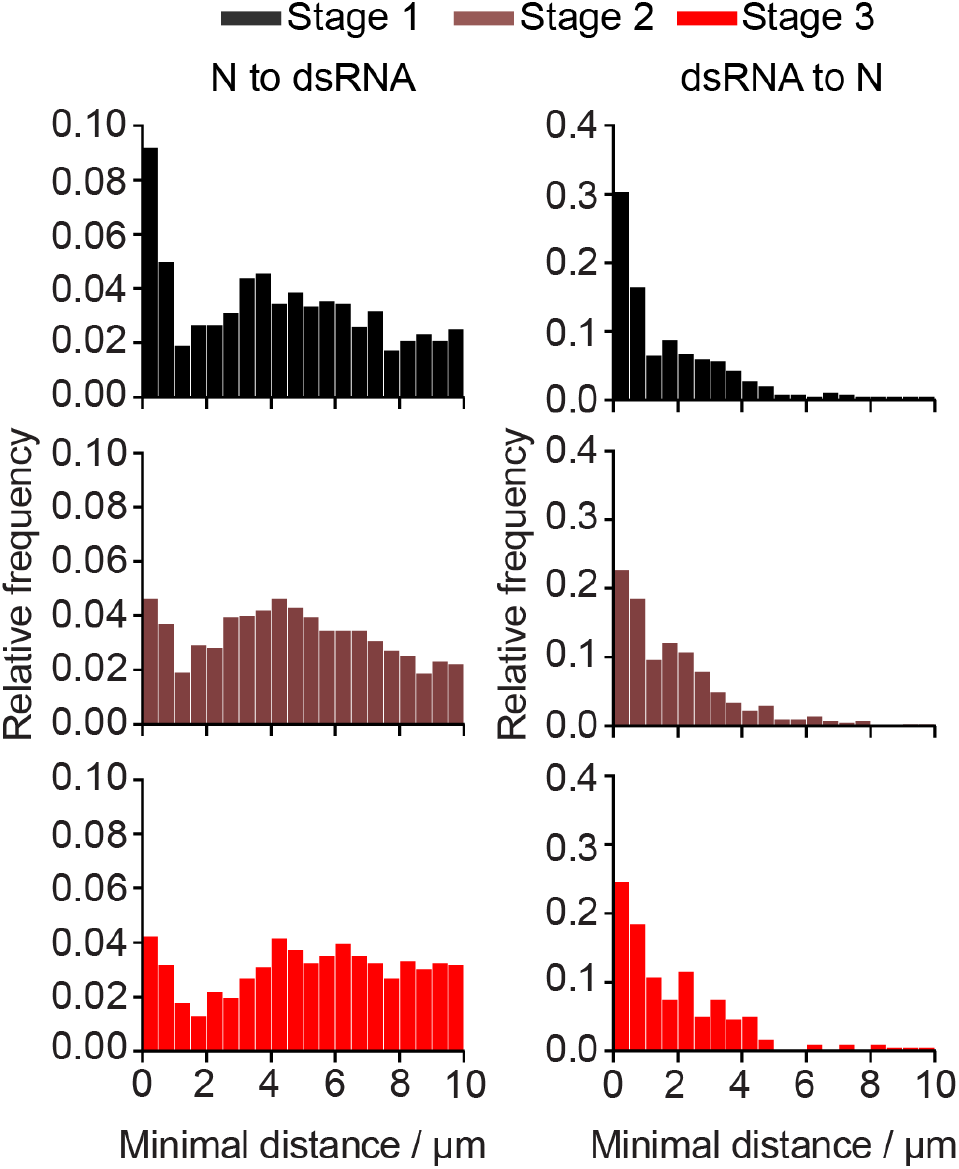
Minimum distance analysis shows that the fraction of closely associated (within <1 μm) N compartments and dsRNA foci decreased during stages 2 and 3.

**Supporting Figure 9:**
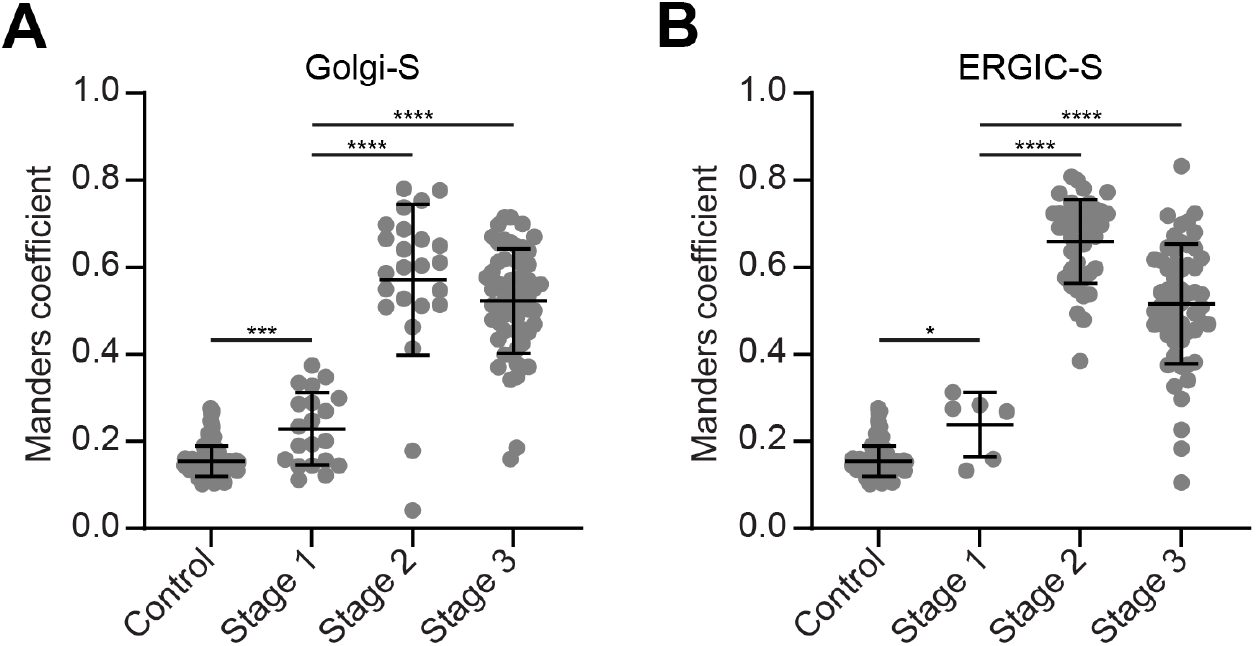
The spike protein (S) of SARS-CoV-2 colocalises with both the Golgi apparatus and the ER to Golgi compartment (ERGIC) in infected Vero cells. A) Colocalisation analysis (determined using the Manders coefficient method) between the Golgi apparatus and S shows partial spatial correlation from the moment S starts to be expressed (stage 2) onwards. B) Colocalisation analysis (determined using the Manders coefficient method) between the ERGIC and S shows partial spatial correlation from the moment S starts to be expressed (stage 2) onwards. At least 20 cells were analysed for each infection stage and organelle except for ERGIC-stage 1, where only six cells were analysed. Significance was tested with an unpaired t-test with Welch’s correction for unequal standard deviations.

**Supporting Figure 10:**
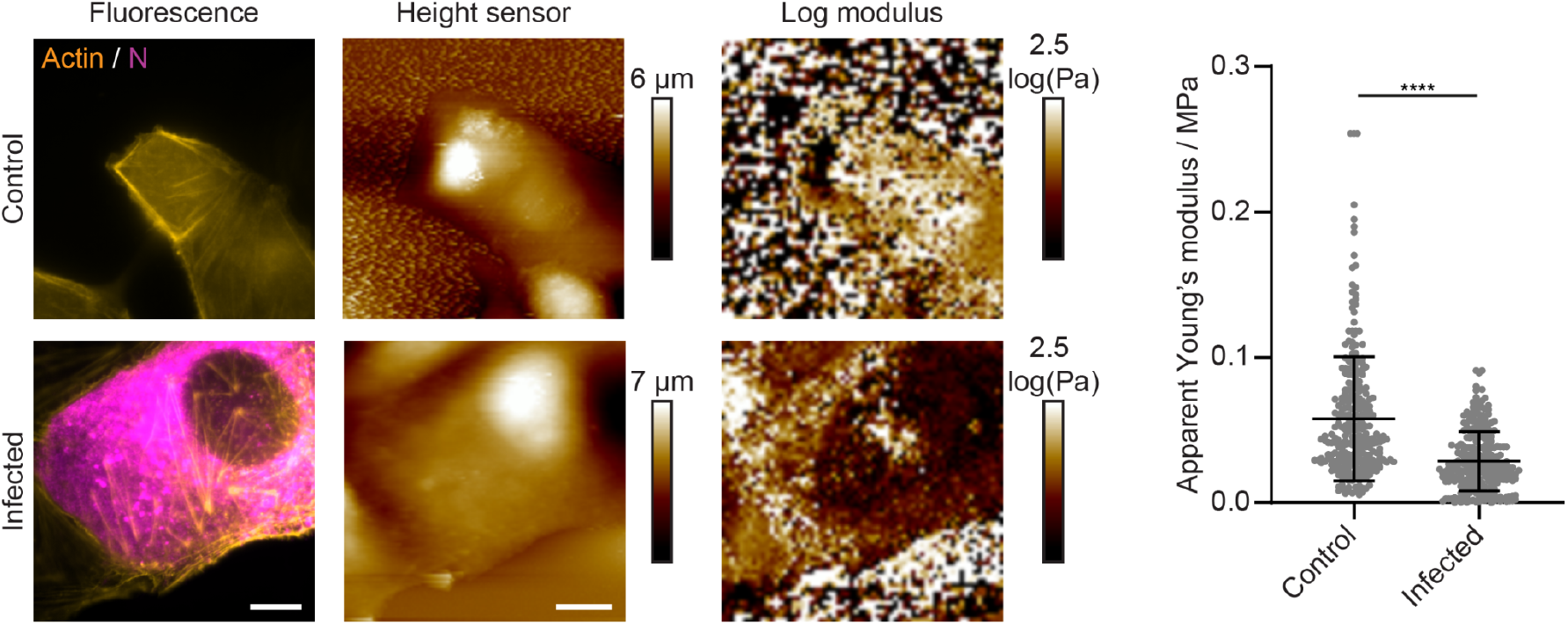
The apparent Young’s modulus is reduced in infected cells which indicates a softening of the probed cell surface compared to non-infected cells. Correlative fluorescence and atomic force microscopy was performed on SARS-CoV-2 infected cells fixed at 12 hpi. Staining of the actin cytoskeleton and of the SARS-CoV-2 nucleocapsid protein (N) was used to distinguish between infected and non-infected cells. The Young’s modulus was obtained by fitting the individual force curves in each picture. Each data point in the graph represents one force curve; for each condition, force curves were obtained from 8 independent cells. Scale bars: 10 μm. Significance was tested using a Mann-Whitney test.

**Supporting Video 1:**
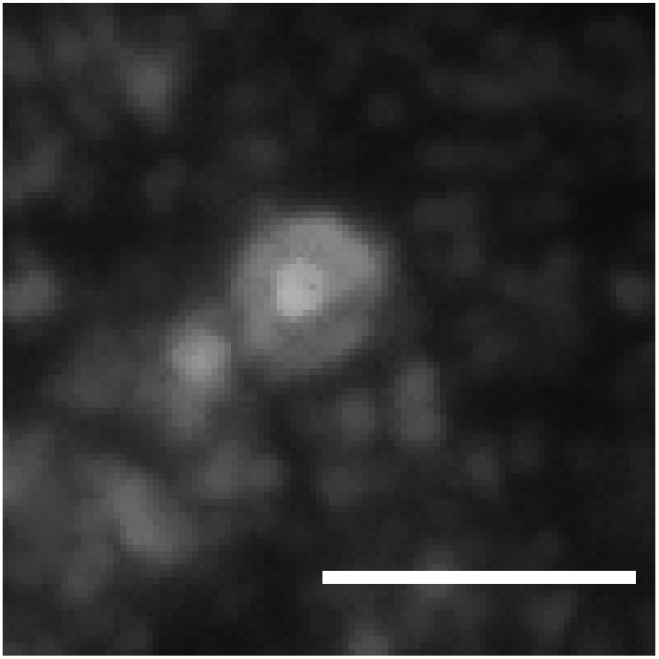
The video shows a single N protein compartment consisting of two layers. SARS-CoV-2 infected Vero cells were fixed, immunostained, expanded and imaged on a light sheet microscope. Scale bar 1.2 μm (taking into account an expansion factor of 4.2).

**Supporting Video 2:**
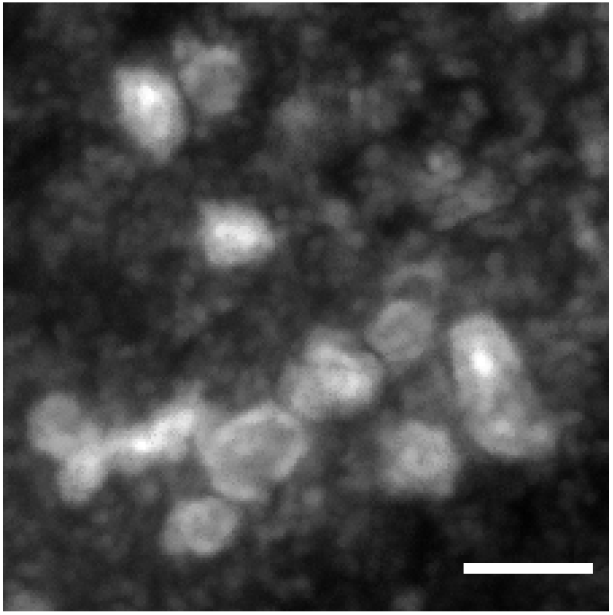
Multiple N protein compartments, partially fused and consisting of two layers. SARS-CoV-2 infected Vero cells were fixed, immunostained, expanded and imaged on a light sheet microscope. Scale bar 1.2 μm (taking into account an expansion factor of 4.2).

**Supporting Video 3:**
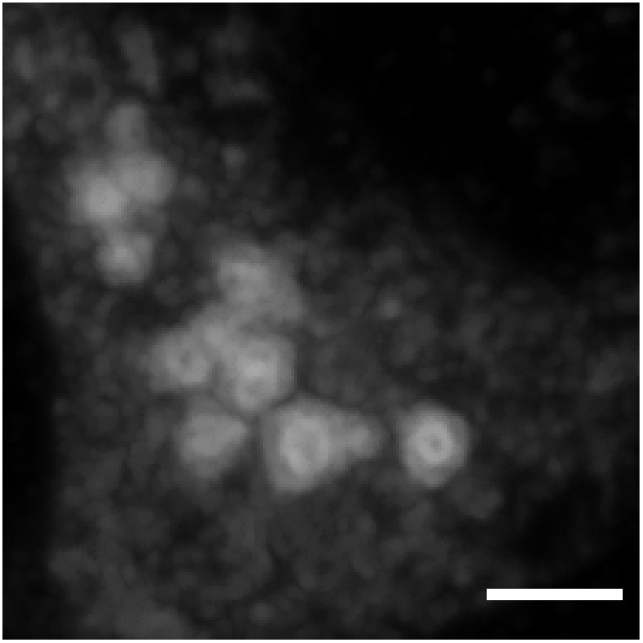
Multiple N protein compartments, partially fused and consisting of two layers. SARS-CoV-2 infected Vero cells were fixed, immunostained, expanded and imaged on a light sheet microscope. Scale bar 1.2 μm (taking into account an expansion factor of 4.2).

**Supporting Video 4:**
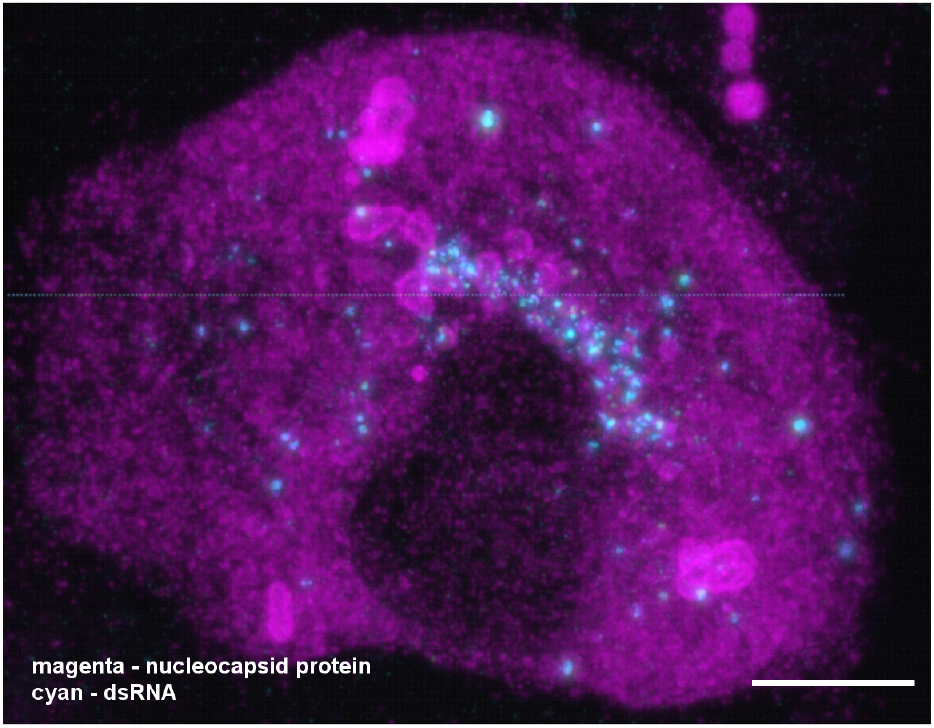
A combination of expansion and light sheet microscopy reveals the interaction between dsRNA (cyan) and the nucleocapsid protein N (magenta) in SARS-CoV-2 infected Vero cells. The video corresponds to Figure 3G in the main manuscript. The majority of dsRNA foci are located in a region immediately adjacent to the nucleus. In addition, most N compartments contain dsRNA foci sitting in the layers of the compartments. Scale bar 5 μm (taking into account an expansion factor of 4.2).

**Supporting Table 1:** Optimised fixation and permeabilization conditions for the immunostaining of the structures imaged in this work. A tick represents a working combination, while a cross represents a non-working combination.

|  | Formaldehyde |  | Glyoxal |  |
| --- | --- | --- | --- | --- |
|  | Saponin | Triton X-100 | Saponin | Triton X-100 |
| SARS nucleocapsid protein | ✓ | ✓ | ✓ | ✓ |
| SARS Spike protein | ✓ | ✓ | ✓ | ✓ |
| SARS Membrane protein | ✓ | ✓ | ✓ | ✓ |
| SARS Envelope protein | ✓ | ✓ | ✓ | ✓ |
| Double-stranded RNA | ✗ | ✓ | ✗ | ✓ |
| ER (calnexin) | ✗ | ✗ | ✓ | ✓ |
| Golgi (GM130) | ✓ | ✓ | ✗ | ✗ |
| ERGIC (LMAN1) | ✗ | ✗ | ✓ | ✗ |
| Lysosomes (LAMP1) | ✓ | ✗ | ✓ | ✗ |
| Microtubules (beta-tubulin) | ✓ | ✓ | ✓ | ✓ |
| Microfilaments (actin) | ✗ | ✗ | ✗ | ✓ |

**Supporting Table 2:** Summary of fixation and permeabilization conditions for all the samples shown in the figures of the manuscript.

| Figure | Fixation | Permeabilization |
| --- | --- | --- |
| Fig. 1A, stage 1 and 3 | Glyoxal | Saponin |
| Fig. 1A, stage 2 | Formaldehyde | Triton |
| Fig. 2A, 2B, 2C, 2E, 2F, 2G | Glyoxal | Saponin |
| Fig. 3A, 3C | Glyoxal | Saponin |
| Fig. 3B | Formaldehyde | Triton |
| Fig. 4A | Formaldehyde | Saponin |
| Fig. 4B, 4C | Glyoxal | Saponin |
| Fig. 5B, 5C | Glyoxal | Saponin |
| Sup. Fig. 2 | Glyoxal | Saponin |
| Sup. Fig. 4 (top row) | Glyoxal | Saponin |
| Sup. Fig. 4 (top row) | Formaldehyde | Triton |
| Sup. Fig. 5B, 5C | Glyoxal | Saponin |
| Sup. Fig. 9 | Formaldehyde | Triton |

